# Engineering kidney developmental trajectory using culture boundary conditions

**DOI:** 10.1101/2024.12.10.627798

**Authors:** Aria Zheyuan Huang, Louis S. Prahl, Karen Xu, Robert L. Mauck, Alex J. Hughes

## Abstract

Kidney explant cultures are traditionally carried out at air-liquid interfaces, which disrupts 3D tissue structure and limits interpretation of developmental data. To overcome this limitation, we developed a 3D culture technique using hydrogel embedding to capture morphogenesis in real time. We show that 3D culture better approximates *in vivo*-like niche spacing and dynamic tubule tip rearrangement, as well as *in vivo*-like presentation of branching defects under perturbations to glial cell-derived neurotrophic factor (GDNF)-<u>RE</u>arranged during <u>T</u>ransfection (RET) tyrosine kinase signaling. We find that the concentration of the embedding matrix influences the number of nephrons per ureteric bud (UB) tip and the spacing between tips. To isolate the effect of specific material properties on explant development, we introduce engineered acrylated hyaluronic acid hydrogels that allow independent tuning of stiffness and adhesion. We find that sufficient stiffness and adhesion are both required to maintain kidney shape. Matrix stiffness has a “Goldilocks effect” on the nephron per UB tip balance centered at ∼2 kPa, while higher matrix adhesion increases nephron per UB tip ratio. Our technique captures large-scale, *in vivo*-like tissue morphogenesis in 3D, providing a platform suited to contrasting normal and congenital disease contexts. Moreover, understanding the impact of boundary condition mechanics on kidney development benefits fundamental renal research and advances the engineering of next-generation kidney replacement tissues.

## Introduction

Branching morphogenesis is a common developmental program for epithelial organs to acquire a tree-like structure. This massively parallelizes their function because branch tips become sites that facilitate organ-specific functions, such as secretion, gas exchange, or filtration^1^. In the embryonic kidney, ureteric bud (UB) progenitor cells exchange reciprocal signals with surrounding mesenchyme to build the branched urinary collecting system and the blood-filtering nephrons^2^. Mammalian kidney architecture and the total number of nephrons (nephron endowment) are determined predominantly *in utero*^3^. Disruptions to branching morphogenesis result in congenital anomalies, where aberrant tissue-scale organization can hinder adult renal function and negatively impact health^4,5^. Therefore, it is crucial to understand the factors involved in the spatial patterning of kidney components during development.

Explant culture techniques enable real-time imaging of a range of processes including branching morphogenesis and nephron formation^6–8^. Branching morphogenesis persists in mouse kidneys that are explanted sufficiently early in development (before embryonic day (E)14) and placed at an air-liquid interface (ALI) on glass, a transwell membrane, or a metal mesh with filter overlay^9–11^. The failure of explants to develop in suspension has been ascribed to a need for appropriate transport of oxygen or other nutrients ^11^. Despite this hypothesis, explant culture also fails in ‘hanging drop’ format, which is thought to have little mass transport limitation^11^. An alternative view arose from the observation that ALI culture is compromised in the presence of a surfactant, suggesting that either mechanical stresses imparted by surface tension and/or kidney-substrate adhesion improve explant morphogenesis^11^. Supporting this, other culture approaches that compress the kidney between material interfaces are similarly successful^8,12^.

However, ALI-based explant culture presents challenges for questions related to mechanics. While genetic and biochemical regulators of kidney branching have been extensively studied, mechanics is of emerging interest. Tissue mechanics are a key contributor to branching morphogenesis in organs including the lung, salivary gland, mammary gland, and pancreas^13–17^. In the kidney, we previously found that limited organ surface area places constraints on the geometry of the ureteric tubule tree such that branching events generate local mechanical stresses on the surrounding nephrogenic mesenchyme^18,19^. Current ALI cultures flatten and distort the cortico-medullary structure of the kidney at an air-liquid interface (ALI), which introduces developmental artifacts^20^. Rosines et al. reported sustained kidney development by embedding explants in 3D matrigel, collagen I, or collagen IV matrices at an ALI^21^. However, the relationship between matrix properties and developmental outcomes has not been closely studied in this context.

Although little is known about the effect of matrix properties on *in vivo* kidney development, explants acquire divergent phenotypes when extracellular matrix (ECM) proteins are added to culture media vs. when bound to the culture substrate in the ALI format^22^. Matrix mechanical properties also influence kidney morphogenesis and cell decision-making in organoid models. For example, hydrogel stiffness and viscoelasticity have been found to affect kidney organoid differentiation, final morphology, and proportions of nephron segments^23–25^. *In vivo*, the kidney is embedded within the urogenital system, bounded by the renal interstitium, capsule, renal fascia, and surrounding organs (**Fig. 1A,B**)^26^. FOXD1+ stromal progenitors form both the capsule and the outermost stroma adjacent to nephron progenitor niches (the capsule is physically distinct after ∼E14)^27^. Genetic ablation of these cells leads to severe defects which are thought to be due to signaling dysregulation^27–30^. However, disruption of the capsule may also play a role through mechanics. We sought to develop a live imaging-compatible 3D culture system that would advance the study of *in vivo*-like branching morphogenesis of the kidney and resolve the influence of mechanical boundary conditions.

**Fig. 1:**
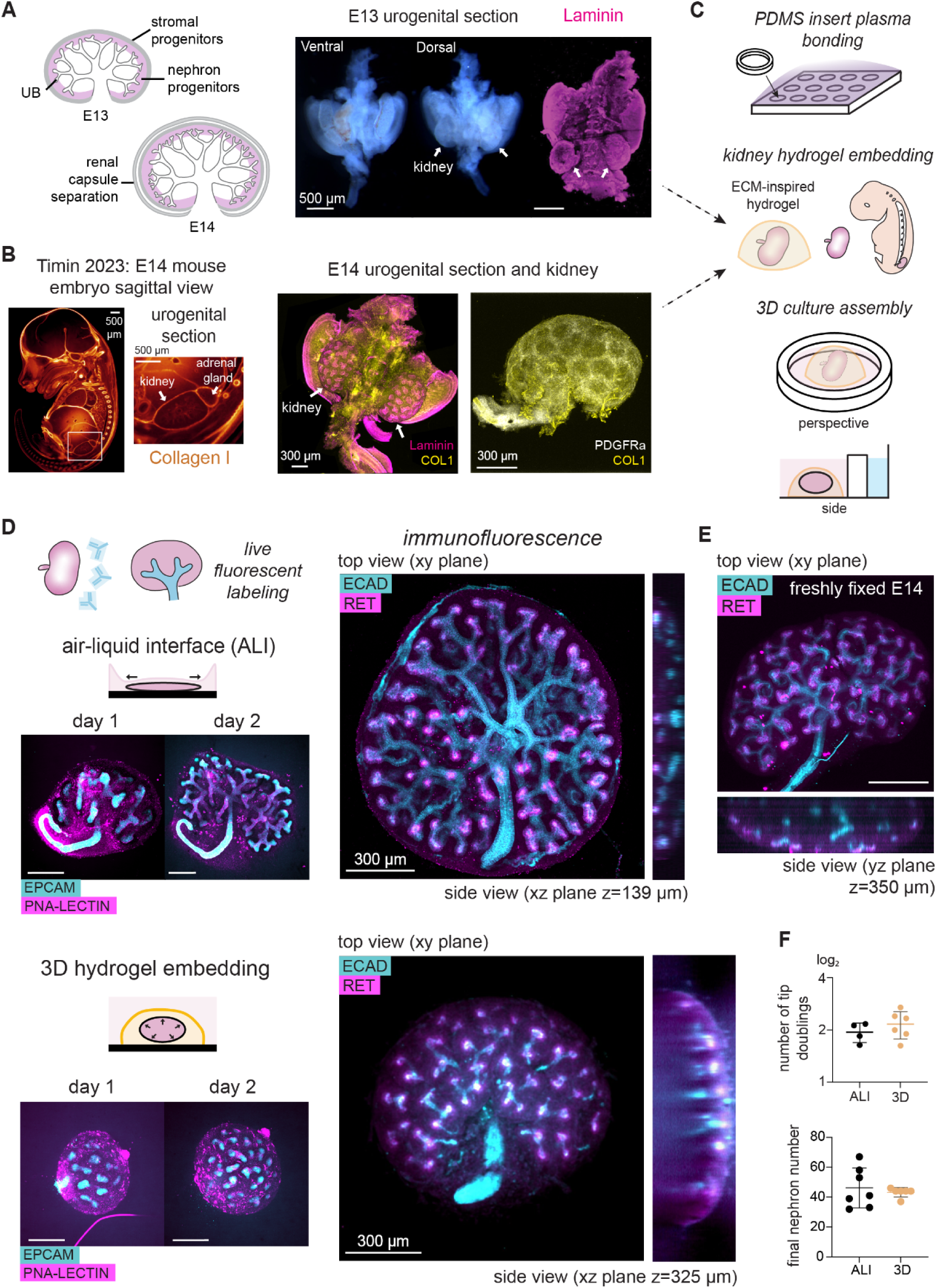
ECM-inspired hydrogel encapsulation supports 3D kidney explant culture. (**A**) *Left*, Schematic of kidney development and renal capsule specification, forming a distinct tissue layer at ∼E14^27,39^. *Right*, Transmitted light and immunofluorescence images of the E13 mouse embryo urogenital tissue region, indicating position of the kidneys (white arrows). (**B**) *Left*, E14 whole embryo fluorescence image of Fast Green-stained collagen 1 (produced from image data associated with ref. ^33^). White arrows pointing toward the kidney and the adrenal gland. Note intense staining in capsule/fascia layers surrounding the kidney. *Middle*, Immunofluorescence image of E14 urogenital tissue region stained for laminin and collagen 1, indicating position of the kidneys (white arrows). *Right*, Isolated E14 kidney stained for collagen 1 and stroma marker PDGFRa to visualize the organ surface. (**C**) Schematic of 3D hydrogel embedding approach for extended mouse embryonic kidney culture. Explanted kidneys are placed in individual 10 µl hydrogel precursor droplets, which subsequently solidifies. Culture media surrounding the hydrogel+kidney is contained in a PDMS ring, allowing humidification of the well using an outer media reservoir. (**D**) Schematic, live, and immunostained confocal fluorescence images of E13 kidney culture in traditional Sebinger air-liquid interface (ALI) culture format (*top*), vs. C+M hydrogel-embedded culture (*bottom*). *Left*, Time point images of kidneys during culture. Live kidneys are stained with fluorescently-labeled antibodies to mark the UB (EPCAM) and the UB basement membrane (PNA-lectin). *Right*, Immunofluorescence images of the top view and side view of kidneys after 3 days of culture in ALI (*top*) and 3D hydrogel (*bottom*) formats. Kidneys stained for epithelial marker E-cadherin (ECAD) and UB tip marker RET. (**E**) Similar immunofluorescence images to (**D**) but for size-matched freshly dissected E14 mouse kidney. (**F**) *Top*, Plot of log_2_(final / initial ureteric bud tip number) vs. culture format after 2 days of culture mean ± S.D.; *n* = 4 and 6 kidneys from the same E13 litter for ALI and 3D culture methods, respectively). *Bottom*, Plot of final nephron number in two culture formats after 3 days. (mean ± S.D.; Top, *n* = 7 and 6 kidneys from the 2 E13 litters for ALI and 3D culture methods).

Inspired by the *in vivo* urogenital environment, we pursued kidney embedding in 3D hydrogel domes to mimic its native encapsulation. We demonstrate that embryonic kidneys develop and acquire more *in vivo*-like morphologies in 3D culture relative to ALI culture. We investigate 3D kidney branching morphogenesis in real time, confirming predictions of tip rotation dynamics and tip-tip spacing inferred from physics-based modeling and fixed kidneys at different ages. Relative to ALI cultures, 3D cultures more faithfully captured *in vivo* effects of perturbing the GDNF-RET pathway, which drives UB branching. We next show that overall kidney shape and nephrogenesis efficiency are influenced by material properties in both ECM-derived and engineered 3D matrices.

## Results

### 3D hydrogel embedding supports kidney explant culture and morphogenesis

We first designed a 3D culture system that embeds mouse embryonic kidneys in hydrogel domes. We began with an ECM-derived hydrogel consisting of 1:1 collagen I and reconstituted basement membrane (Matrigel). Collagen I is a structural protein that provides tensile strength in the ECM^31,32^. Whole E14 embryo staining revealed that collagen I-rich ECM envelops the developing kidney *in situ*^33^ (**Fig. 1B**). Matrigel mainly comprises laminin and collagen IV, two ECM components abundant in embryonic kidneys (**Fig. 1A,B**)^34,35^. Mixing 1 mg/ml collagen I and Matrigel (C+M) at a 1:1 ratio results in 0.5 mg/ml C+M gels that adhered to glass culture substrates, gelled consistently, and had sufficient mechanical integrity to support a consistent 3D geometry during culture over at least 3 days (**Fig. S1**). To contain overlying media, we fabricated and plasma-bonded polydimethylsiloxane (PDMS) rings to coverglass plates to separate the inner space for explant culture and the outer space for humidity control (**Fig. 1C**).

Next, we aimed to apply a labeling strategy compatible with time-lapse fluorescence microscopy. Inspired by studies in lung explant systems^36^, we labeled live kidneys with fluorescently-conjugated EPCAM antibody and PNA lectin (both stains for the ureteric bud, **Fig. 1D**). This approach allowed us to capture time-lapse movies of 3D branching morphogenesis using spinning disk confocal microscopy (**Fig. 1D**, **Movie S1**). By quantifying UB tip numbers from time-lapse data, we found that tip duplication rates were equivalent between 3D and ALI cultures (**Fig. 1E**). After ∼3 days (64 hours), we compared cultures in 3D C+M to those in ALI by immunofluorescence. While branching morphogenesis proceeded in both cultures, kidneys in ALI culture flattened to ∼140 µm in height. Kidneys in C+M culture better maintained their native thickness at ∼325 µm in height, as compared to size-matched E14 kidneys with thickness at ∼350 µm (**Fig. 1D,E**). We noted that RET+ UB tips were localized at the organ surface in 3D, while some tips were found internal to the kidney in ALI culture. This is likely due to surface area constraints in flattened ALI cultures, where branching morphogenesis creates more tips than can be geometrically accommodated at the organ periphery^18^. Tip localization at the organ surface is a key feature of kidney morphogenesis and is likely necessary to properly localize nephron condensation events to the (outer) cortex relative to the (inner) medulla, contributing to appropriate corticomedullary zonation in the adult organ^37,38^. We then assessed the number of UB tips and nephrons and found them to be comparable in both culture formats (**Fig. 1F, Fig. S2**). Together, these data indicate that kidney embedding in C+M hydrogels retained live imaging capabilities and similar UB tip and nephron numbers to ALI culture, but better approximated *in vivo*-like tissue thickness and localization of UB tips to the organ periphery.

### 3D culture reveals distinct morphogenesis patterns that are not present in ALI culture

We next focused on live dynamics of tip branching. We found that tip branching proceeded in 3D in embedded cultures, with tips bifurcating and advancing in *xy* and *z* (**Fig. 2A**). During branching *in vivo*, UB tips become more crowded at the kidney surface, eventually forming semi-crystalline packing after ∼E15 in mice and week ∼11 in humans^18,19^. We wondered which culture approach better approximated this close packing of UB tips. We quantified the distance between non-sibling tips as a metric for tubule packing. E13 kidneys cultured for ∼3 days in C+M hydrogels and size-matched E14 kidneys had similar UB tip-tip distances, while tips in ALI cultures were further away from their neighbors (**Fig. 2B**). These data indicate that UB tips more closely adopt *in vivo-like* packing in 3D culture relative to ALI culture.

**Fig. 2:**
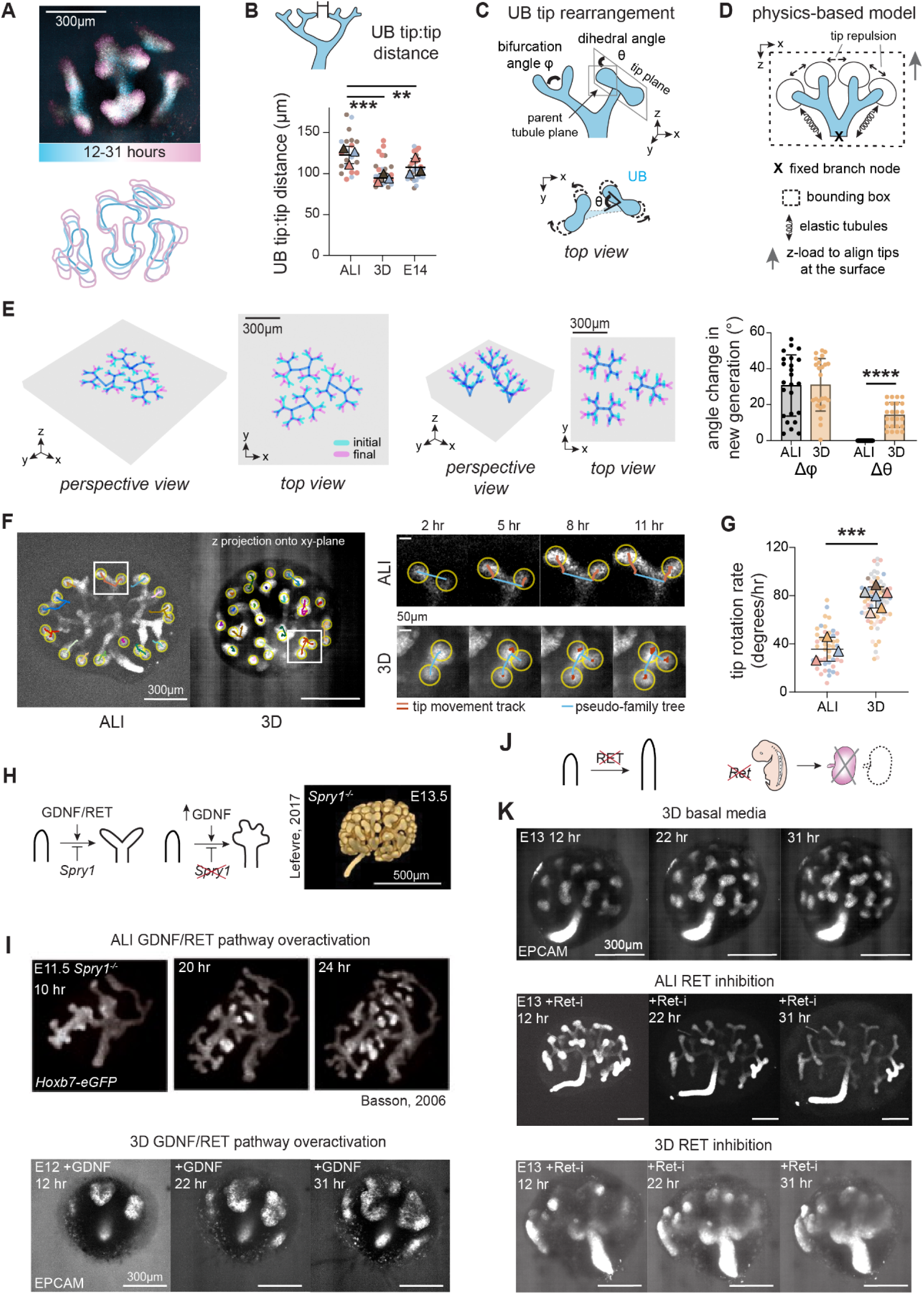
3D culture models kidney morphogenesis dynamics under normal conditions and signaling perturbations. (**A**) *Top*, A representative average immunofluorescence projection of E12 embryonic kidney in 3D 0.5C+M hydrogel culture, color-coded by culture time. *Bottom*, Manually annotated outlines of ureteric bud at each time point, highlighting the branching process. (**B**) Schematic and plot of E13 kidney UB branch tip distance after culture in ALI or 3D formats for 3 days, compared to the same measurement for dissected and immediately fixed size-matched E14 kidney controls (mean ± S.D., n = 21, 24 and 34 tip pairs for ALI, 3D, and E14 categories across 3 kidneys each, pooled across 3 independent litters, respectively). Tip distances were annotated for tips with a branching relationship at least two generations apart (non-sibling or cousin tips). One-way ANOVA with Tukey’s test, p=0.0002 and 0.0016 for between ALI and 3D culture and between ALI and E14. (**C**) Schematic of tip rotation during branching by changing its dihedral angle. (**D**) Schematic of physics-based model definition. (**E**) Simulation and quantification of branching of UB trees under physical constraints that model 2D ALI (*Left*) vs. 3D hydrogel (*Middle*) culture formats. *Left and middle*, perspective and top views of the UB branch families before and after energy minimization within the bounding box (gray). *Right*, quantification of branching angle change before and after energy minimization (mean ± S.D., *n* = 24 and for both 𝜙 and θ, p < 0.0001). (**F**) Live fluorescence images showing tracking of E13 kidney UB tip positions over 11 hr of culture in ALI vs. 3D hydrogel cultures. Insets show tracked tips (yellow circles) and tip position tracks (red/brown lines) for the indicated time points. (**G**) Plot of track rotation velocity in ALI vs. 3D culture (mean ± S.D., *n* = 45 from 3 E12 and 1 E13 kidneys for ALI culture and 75 tips from 4 E12 and 2 E13 kidneys for 3D culture, pooled across 2 and 3 litters, respectively, p<0.0001). (**H**) *Left*, Schematic of GDNF-RET signaling and GDNF overactivation. Summarizing effects of *Spry1^-/-^* (supernumerary tips). *Right*, Top: 3D optical projection tomography reconstruction of *Spry1^-/-^* kidney UB branch pattern at E13.5, reproduced from Lefevre *et al*.^20^. Note enlarged UB tips. Bottom: Schematic of *Ret* knockout effects leading to renal agenesis. (**I**)*Top*, Live fluorescence images of E11.5 *Hoxb7-eGFP;Spry1^-/-^* kidney branching in ALI culture, reproduced from Basson *et al.*^44^. UB is marked by the *Hoxb7-eGFP* reporter. *Bottom*, Live fluorescence projections of E12 kidneys at the indicated time points in 3D culture in the presence of 100 ng/ml recombinant GDNF. (**J**) Schematic of RET downregulation effects in culture and *in vivo*. (**K**) Similar images to (**H**) but for E13 culture in basal media in 3D C+M gel (*top*), with 100nM of the RET inhibitor selpercatinib in ALI transwell (*middle*), and with RET inhibitor in 3D gel (*bottom*). Unpaired two-tailed t-test, *p < 0.05, **p < 0.01, ***p < 0.001, ****p < 0.0001.

We speculated that tissue flattening in ALI culture could account for changes in tip packing by creating branching artifacts that are not representative of *in vivo* organ growth. Retrospective studies of kidneys dissected at different developmental ages have inferred that UB tips rotate and reorganize at the organ surface throughout development^18,19,40,41^ (**Fig. 2C**). Our prior physics-based model also revealed that this tip organization *in vivo* could be explained by physical packing rules in 3D^18^. The model performs energy minimization to predict elastic tubule positions within a 3D bounding box and repulsion forces between tips that are inversely proportional to their distances^18^ (**Fig. 2D)**. We applied the same model to simulate tip movements in ALI and 3D cultures. To provide a quantitative comparison of tip rearrangement due to packing, we defined a bifurcation angle 𝜙 and a ‘dihedral’ tip rotation angle θ, similar to ref. ^41^. We previously quantified 𝜙, which becomes smaller over developmental time *in vivo*^18,42^. However, our interest here was to investigate the shorter timescale variation in θ caused by tips accommodating each other due to branching amidst crowding (**Fig. 2C**). We simulated initial tree geometries for bounding box constraints that were consistent with ALI and 3D culture formats, allowing tubules to adopt energetically favorable positions (see **Materials & Methods**). We then added a new pair of daughter tubules to each existing branch, and measured 𝜙 and θ before and after a second simulation step (**Fig. 2E, Fig. S3**). We found that the change in θ was significantly higher in the model case simulating 3D culture relative to that simulating ALI, while the change in 𝜙 was similar between the two. This predicts that kidney flattening in ALI culture, constraining tubules to the same plane, leads to a loss of the dihedral rotational degree of freedom.

To validate the geometric model predictions in our experimental system, we projected time-lapse movies of UB tip movements from ALI and 3D cultured onto the *xy*-plane and annotated UB branching tracks (**Fig. 2F**). Our analysis revealed that branching tips in ALI culture primarily elongate linearly outward in the radial direction from the center of the kidney as indicated by small rates of change in the directions of their elongation tracks. On the contrary, tips in 3D culture exhibited higher rates of change in their elongation directions due to ongoing repositioning at the explant surface (**Fig. 2G, Movie S2,S3**). This suggests that the linear branching progression in ALI is likely an artifact generated by their limited mobility in the *z*-direction, essentially confining tip movement/rearrangement to a 2D-like environment. This result validates packing-based tip repositioning that had previously only been inferred from fixed kidneys and geometric modeling. Taken together, the data emphasize that while ALI culture constrains branching and packing to the same *xy* plane, 3D hydrogel culture retains distinct tip branching and packing planes as *in vivo* (*xz*/*yz* and *xy*, respectively). This restricts ALI cultures to the study of processes operating at the length scale of individual tips and nephron condensation events, but not to those operating at the scale of groups of tips or the whole organ, including those giving rise to many kidney defects.

Such a capability for capturing larger-scale 3D developmental dynamics in real-time would benefit the study of congenital kidney defects, where the nature or timing of the fault along the developmental trajectory is often obscured. In this vein we applied our 3D culture method to the glial cell-derived neurotrophic factor (GDNF)-<u>RE</u>arranged during <u>T</u>ransfection (RET) pathway, which is central to kidney branching. Prior work has established that increased GDNF activation by either ectopic delivery of GDNF or knockout of a negative feedback regulator *Spry1* leads to more closely spaced branch buds and dilated tips that have been observed to merge in ALI culture^43–45^ (**Fig. 2H**). We added GDNF to 3D E12 mouse kidney cultures and compared their morphology to published *Spry1^-/-^* mutants^20,43^ (**Fig. 2I**). We noted that the morphology of the mutant is different in freshly dissected kidneys compared to those explanted and cultured at the ALI. Specifically, *Spry1^-/-^* kidneys in ALI culture show clustered branches with many ‘buried’ tips. However, freshly dissected *Spry1^-/-^* kidneys primarily have dilated UB tips rather than clustered branches. Our +GDNF 3D cultures had dilated tips, more similar to the *in vivo* presentation of the *Spry1^-/-^* mutant compared to its presentation in ALI culture (**Fig. 2H, Movie S4**).

On the other end of the GDNF-RET axis, loss of RET signaling is detrimental to kidney development, causing renal hypoplasia and agenesis^46–48^. GDNF-RET inhibition by adding either a GDNF-neutralizing antibody or a MEK inhibitor to block RET-downstream signaling abolishes tip bifurcation in culture, leading to fewer tips in total but retaining persistent tubule elongation^45,49^ (**Fig. 2J**). We confirmed that UB tips in E13 kidneys cultured in ALI format with the selective RET inhibitor (RET-i) selpercatinib^50^ continued to elongate but did not bifurcate, relative to those in littermate control kidneys cultured without RET-i (**Fig. 2K**). Surprisingly, UB tubules buckled and folded back on themselves while elongating under RET-i conditions in 3D culture, perhaps reflecting the reduced kidney spreading area-to-volume ratio in 3D culture relative to ALI (**Fig. 2K, Movie S5**). This morphology resembles a characteristic of renal hypoplasia resulting from retinoic acid signaling inhibition, an upstream regulator of RET^51^. Together, our experiments with GDNF-RET perturbations indicate that 3D hydrogel culture may more faithfully model morphology changes in response to developmentally relevant defects compared to ALI culture.

### ECM-derived matrix composition impacts final kidney morphology

In prior work, Sebinger et al. found that the composition and concentration of ECM proteins affect kidney development when presented at the interface between kidney and substrate in ALI cultures^22^. The implication here is that interfacial properties at the boundary of the kidney can affect relatively distant cell and tissue-level behaviors in the ureteric bud and nephrogenic niches at their tips. We hypothesized that one category of properties here could be mechanical, and set out to test this using our 3D culture technique. Explants cultured in fluorescently pre-labeled collagen 1-containing C+M hydrogels pulled the ECM inward over the culture period, indicating that kidneys actively exert a contractile force on their embedding matrix normal to the interface between the two (**Movie S6**). More generally, we can conceive of a force balance at each point along the interface between forces due to growth, matrix elasticity, tissue contractility, and shear stress due to cell traction (**Fig. 3A)**. Such stresses may feed into morphogenesis through effects on cell migration, differentiation, and proliferation^52,53^.

**Fig. 3:**
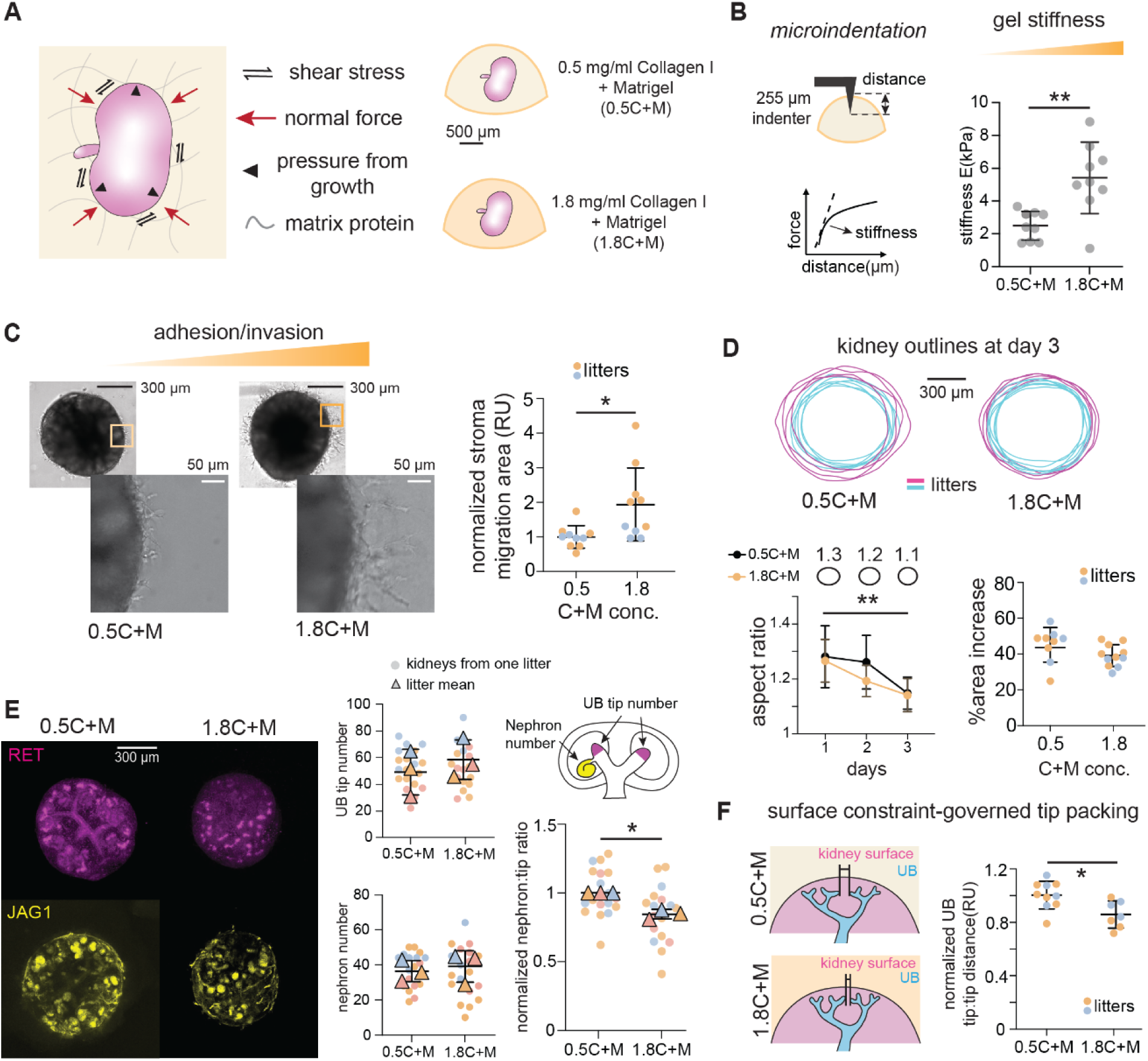
3D Matrigel-collagen hydrogels allow global kidney rounding and affect nephron:UB tip ratio upon extended culture. (**A**) *Left*, Schematic of 0.5C+M and 1.8C+M hydrogel embedding conditions. *Right*, model of force balance determining kidney size and shape through interactions between kidney growth, and stiffness and adhesion properties of the surrounding microenvironment. (**B**) *Left*, schematic of hydrogel microindentation and inference of elastic stiffness. *Right*, Plot of hydrogel stiffness vs. composition. Measurements taken one day after swelling hydrogels in culture media to mimic the culture environment (mean ± S.D.; *n* = 9 measurements from 3 technical replicate gels for both 0.5C+M and 1.8C+M conditions; p = 0.0036). *E*, Young’s modulus. (**C**) Phase contrast micrographs of E13 kidneys after 3 days of culture in 0.5C+M (*left*) and 1.8C+M (*middle*) hydrogels, highlighting difference in interaction and invasion of presumptive stromal progenitor cells into the gel. *Right*, Plot of normalized segmented area of stromal invasion at kidney midplanes (mean ± S.D.; *n* = 10 kidneys pooled from 2 litters for both 0.5C+M and 1.8C+M culture conditions; p = 0.0152). (**D**) Top: Outlines of kidneys cultured in 0.5C+M and 1.8C+M hydrogels after 3 days for two litters (cyan, magenta). Bottom: Plots of kidney outline aspect ratio and % growth in midplane area after 3 days of culture (mean ± S.D.; *n* = 9 and 11 kidneys pooled from 2 litters for both 0.5C+M and 1.8C+M culture conditions). *Left*: two-way ANOVA with Tukey’s test, p=0.0015 and 0.0013 for 0.5C+M and 1.8C+M; *right*: unpaired two-tailed t-test. (**E**) *Left*, Immunofluorescence projections of kidneys after 3 days of culture in 0.5C+M and 1.8C+M hydrogels, stained for UB (RET) and early nephron (JAG1) markers. *Right*, Plots of UB and nephron number and normalized nephron:UB tip ratio vs. hydrogel type after 3 days of culture (mean ± S.D.; n = 19 kidneys pooled across 3 litters for both 0.5C+M and 1.8C+M culture conditions; p = 0.0149). (**F**) Schematic and plot of UB tip-tip distance vs. hydrogel type for non-sibling tips. Plot quantifies E13 kidney UB branch tip distance after culture in 0.5C+M and 1.8C+M for 3 days. Average distance per kidney is plotted with colored dots indicating litter origin (mean ± S.D., *n* = 54 and 37 tip pairs for 0.5C+M and 1.8C+M from 10 and 7 kidneys each, pooled from the same 2 litters, respectively; p = 0.0117). Statistics in (**B**), (**C**), (**E**), and (**F**) are unpaired two-tailed t-tests. All normalization data are normalized to the mean of the 0.5C+M condition from the corresponding litter. RU, relative unit. *p < 0.05, **p < 0.01, ***p < 0.001, ****p < 0.0001.

We expected that the change in collagen concentration would simultaneously affect several material properties including stiffness and adhesion. To begin assaying this, we cultured kidneys in Matrigel with 1.8 mg/ml collagen (1.8C+M) hydrogels and compared outcomes with those cultured in the Matrigel with 0.5 mg/ml collagen (0.5C+M) hydrogels used thus far for live imaging experiments (**Fig. 3A**). The concentration of collagen I gels has been reported to correlate with their stiffness^31,32^, predicting that the same would hold in collagen I-Matrigel composite gels. Indeed, we performed microindentation and confirmed that 1.8C+M hydrogels are stiffer than 0.5C+M hydrogels (**Fig. 3B, Fig. S4**). Aside from stiffness contributions, collagen I harbors receptor-binding sites that promote cell adhesion^54,55^. Therefore, we expected that the adhesion of explants to the surrounding matrix would be higher with the increase in collagen concentration. Migration of stromal cells at the explant surface is commonly seen in both ALI and 3D explant cultures (**Movie S7,S8**). Explants show poor development when cultured in non-adherent conditions where stroma-matrix interaction is absent (**Fig. S5**). In C+M hydrogels, we observed different stromal cell migration patterns in the two collagen concentrations. Stromal cell migration away from the explant was higher in 1.8C+M gels by day 3, as quantified by the relative stromal cell area beyond the explant boundary (**Fig. 3C, Fig. S6**). We interpreted this as being driven by differences in matrix adhesion and/or stiffness. These data indicate that 1.8C+M hydrogels are stiffer and potentially more adhesive to stromal cells than 0.5C+M hydrogels.

To compare developmental outcomes between the two composite hydrogels, we imaged and annotated kidney morphology over 3-day culture. We observed that kidneys converge toward spherical shapes in both conditions, as quantified by progressively decreasing aspect ratio of kidney cross-sectional outlines (**Fig. 3D**). No significant differences were found in the shape or area of kidneys between the two conditions throughout the culture period (**Fig. 3D, Fig. S6**). Despite this, explant studies showed that matrix concentration regulates branching/differentiation outcomes^13,22^. Therefore, we hypothesized that changing the embedding matrix concentration may influence branching morphogenesis and nephrogenesis efficiency. To assess this more closely, we quantified the number of UB tips and early nephrons after culture. Both of these are subject to developmental variability in effective gestational age between littermates at the start of culture^56,57^ (**Fig. 3E, Fig. S7**). To correct for this, we quantified the number of nephrons per UB tip as a proxy for nephrogenesis efficiency in each condition. In developing kidneys *in vivo*, this ratio is thought to be tightly controlled by a balance between nephron progenitor self-renewal vs differentiation^41,58,59^. We found significant decreases in the nephron:UB tip ratio and tip:tip distance in 1.8C+M as compared to 0.5C+M gels (**Fig. 3E,F; Fig. S8**). These data demonstrate that matrix concentration affects UB tip crowding and nephron:UB tip ratio in 3D culture, potentially as a function of either stiffness or adhesion at the kidney-hydrogel boundary.

### Acrylated hyaluronic acid (AHA) hydrogel stiffness and adhesion tune explant geometry and nephrogenesis per UB tip

Since collagen and Matrigel are ECM-derived materials, it is difficult to tune stiffness and adhesion orthogonally. Additionally, the exact compositions of ECM-derived hydrogels are poorly characterized and are subject to batch-batch variability^32^. To delineate the effects of stiffness and adhesion on 3D kidney growth, we turned to engineered biomaterials that offer precise control over composition and material properties. However, certain engineered 3D hydrogels, namely alginate and puramatrix, have failed to support rat kidney development^21^. We first sought to identify a biomaterial suitable for 3D mouse kidney culture. Hyaluronic acid-based hydrogels and gelatin methacryloyl (GelMA) are widely used in tissue engineering and regenerative medicine applications due to their high biocompatibility and tunability^60–62^. In pilot experiments, both C+M and acrylated hyaluronic acid (AHA) hydrogels supported kidney culture, while those in GelMA did not survive, potentially due to free radicals generated during polymerization (**Fig. S9**). We therefore pursued AHA hydrogels consisting of the AHA polymer, matrix metalloproteinase (MMP)-degradable crosslinkers (MMPc), and covalently-conjugated RGD peptides (**Fig. 4A, Fig. S10**). In this system, changes in crosslinker or RGD ligand concentrations can be used to independently tune hydrogel stiffness and cell adhesion, respectively^60^.

**Fig. 4:**
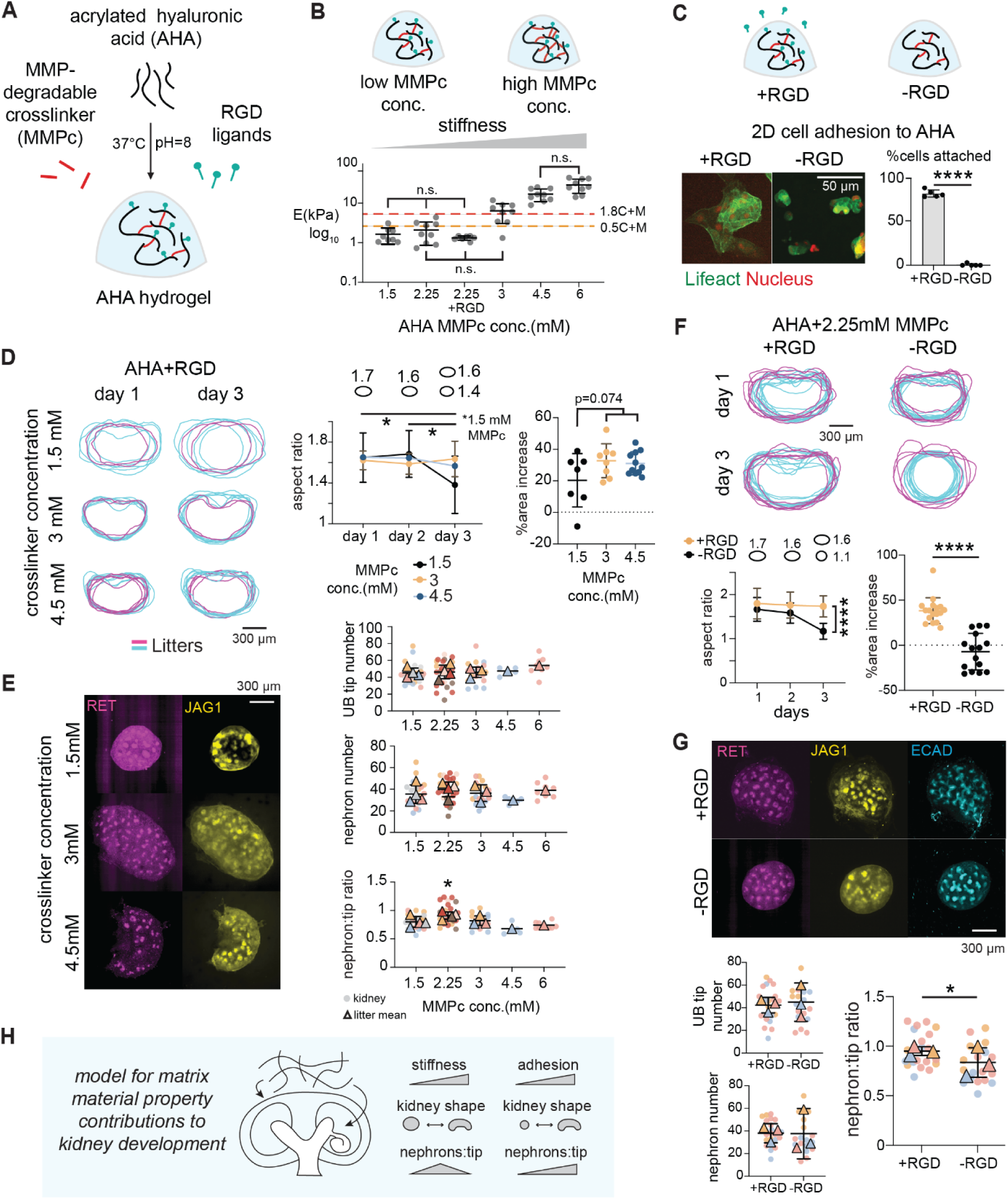
Engineered hydrogel stiffness and adhesion control kidney shape and nephron-tip ratio upon extended culture. (**A**) Schematic of AHA hydrogel crosslinking and conjugation through Michael addition, with MMP-degradable crosslinkers and RGD functional groups enabling controlled stiffness and adhesive properties, respectively. (**B**) *Top*, Sketch of crosslinker concentration effects on AHA gel stiffness. *Bottom*, Plot of hydrogel stiffness by microindentation vs. AHA with MMPc concentrations of 1.5-6 mM (mean ± S.D.; *n* = 8, 8, 7, 9, 9, 9 for AHA with MMPc of 1.5-2.25 mM, 2.25 mM+RGD, 3-6 mM, respectively, measurements taken from 3 technical replicates each, one day after swelling in culture media). All AHA gels are crosslinked without RGD unless noted. Mean stiffnesses of 0.5C+M and 1.8C+M gels are plotted as dotted lines. Groups with no significant difference in pairwise comparison are indicated as n.s.. The rest are significantly different with p<0.05 or less. Full statistical results from Brown-Forsythe ANOVA test included in **Fig. S11**. *E*, Young’s modulus (stiffness). (**C**) *Top*, Sketch of AHA hydrogels +/- RGD functionalization. *Bottom*, confocal immunofluorescence images and plot of MDCK cell attachment to 2D +RGD and -RGD 2.25 mM MMPc AHA hydrogels. Percentage cells attached is defined as % attached cells out of total cells after washing (mean ± S.D.; *n* =5 wells each, unpaired two-tailed t-test, p<0.0001). (**D**) *Left*, Outlines of E13 kidneys cultured in AHA+RGD hydrogels of the indicate crosslinker concentrations after 1 and 3 days for two litters (cyan, magenta). *Right*, Quantifications of the outlines showing kidney aspect ratio and % midplane area increase relative to day 1 for the indicated MMPc concentrations (mean ± S.D.; *n* = 7, 8, and 12 kidneys for 1.5, 3, and 4.5 mM MMPc conditions, respectively, pooled from 2 litters). *Left*, two-way ANOVA with Tukey’s test, p = 0.036 and 0.0265 for between day 1 and 3 and between day 2 and 3 for the 1.5 mM MMPc condition. Sketches above aspect ratio plot illustrate ratios for ellipses equivalent to mean kidney values. *Right*, p = 0.074 from Kruskal-Wallis test. (**E**) *Left*, Confocal immunofluorescence micrographs of kidneys after 3 days of culture in AHA gels of the indicated MMPc concentration, stained for UB tips (RET) and early nephrons (JAG1). *Right*, Plots of UB tip and early nephron number, and ratio of nephrons:tip for AHA gels of the indicated MMPc concentrations (mean ± S.D.; *n* = 21, 24, 18, 4 and 8 kidneys for 1.5-6 mM MMPc conditions, pooled across 7 litters in total). One-way ANOVA with Tukey’s test. p=0.018, 0.045, 0.0085, 0.0093 for comparison between 2.25 mM MMPc and 1.5, 2.25, 3, 4.5 and 6 mM MMPc conditions, respectively. (**F**) Similar kidney outlines, aspect ratio, and % midplane area increase plots as in (**D**), except for +/-RGD functionalization in 2.25 mM MMPc AHA gels (mean ± S.D.; *n* = 17 and 15 kidneys for +/- RGD conditions, respectively, pooled from 2 litters each). *Left*, two-way ANOVA with Tukey’s test, p<0.0001 between +/- RGD. *Right*, unpaired two-tailed t-test, p < 0.0001. (**G**) Similar confocal immunofluorescence images and plots as for (**E**), except for +/-RGD AHA gels (mean ± S.D.; *n* = 20 and 18 kidneys for +/- RGD conditions, respectively, pooled across 3 litters each). One-way ANOVA with Tukey’s test, p=0.0343. (**H**) Schematic summary of figure findings, featuring ‘Goldilocks’ nephron:UB ratio vs. hydrogel stiffness, and increased nephron:UB ratio for adhesive vs. non-adhesive hydrogels. *p < 0.05, **p < 0.01, ***p < 0.001, ****p < 0.0001. Outliers are removed using nonlinear regression (ROUT) method with false discovery rate at 1%.

We first characterized the stiffness of AHA hydrogels over a range of crosslinker concentrations using microindentation. We found that 1.5 mM MMPc was a lower bound for hydrogel integrity over the culture period. *E* scaled with the concentration of MMPc present, validating quantitative control over this parameter. AHA gels with 1.5 and 2.25 mM MMPc and 0.5C+M hydrogels also exhibited similar stiffnesses (**Fig. 4B, Fig. S11**). To confirm orthogonal control over cell adhesion, we plated Madin-Darby canine kidney (MDCK) cells on substrates coated with AHA with 2.25 mM MMPc crosslinker, with and without RGD functionalization. MDCK cells on AHA without RGD did not attach (instead forming spheroids). In contrast, cells successfully adhered and spread on gels containing RGD (**Fig. 4C**). Cell adhesion to AHA therefore depends solely on the presence of the RGD functional groups, which did not affect gel stiffness (**Fig. 4B**). These data indicate orthogonal control over adhesion and stiffness in engineered AHA hydrogels.

To investigate the effect of matrix stiffness on 3D kidney development, we cultured kidneys in AHA+RGD with crosslinkers ranging from 1.5 mM to 6 mM (stiffness between ∼1-40 kPa). Kidneys rounded in the softest hydrogels having 1.5 mM MMPc, similar to those in C+M hydrogels (**Fig. 4D**). However, kidneys grown in AHAs with >= 2.25 mM MMPc maintained their initial “bean-like” shapes (aspect ratio > 1) throughout the culture. We also quantified midplane cross-sectional area increase over time, which showed that kidneys in the softest 1.5 mM AHA gels had a smaller area increase over time relative to those in gels of higher crosslinker concentration (though this effect was not significant, p = 0.074, **Fig. 4D**). Overall, kidneys maintained a more *in vivo*-like elongated shape in AHA+RGD hydrogels. This likely reflects differences in the adhesion profile of the kidney strictly to RGD vs. more diverse ligands in collagen/Matrigel^63,64^ and/or differences in plastic deformation of each gel type in response to tissue-generated stresses that drive rounding. We next quantified UB tips and nephrons in kidneys cultured in AHA+RGD gels with respect to crosslinker concentration (a correlate of stiffness). We noted similar intralitter variability in the number of UB tips and nephrons as seen in C+M cultures (**Fig. 4E**). However, kidneys achieved a significantly higher nephron:UB tip ratio when cultured in AHA with 2.25 mM crosslinker (*E* ∼2 kPa) relative to softer or stiffer ones (14%, 11.3%, 34.3%, and 22.8% increases relative to 1.5, 3, 4.5, and 6 mM MMPc gels, respectively, **Fig. 4E**). These data indicate a “Goldilocks” microenvironmental stiffness that maximizes nephrogenesis efficiency.

We next studied the effect of gel adhesion to kidney morphology and nephrogenesis efficiency in AHA gels, independent of stiffness. Cell adhesion to the ECM is critical for many biological processes including collective cell movements and tissue-building during development^65^. To probe the role of adhesive boundary conditions, we embedded embryonic kidneys in AHA gels with or without RGD functionalization. Similar to MDCK cells, we observed stromal cells interacting with the matrix in adhesive AHA gels but not in those lacking RGD (**Fig. S12**). Quantification of tissue geometry over the culture period revealed a global rounding effect in kidneys cultured without RGD, similar to the phenotype observed in C+M gels and the softest AHA+RGD gels (**Fig. 4F**). In contrast, kidneys in +RGD gels retained their initial elongated shapes throughout culture. Kidney midplane area also radically increased in these gels (by ∼38%), whereas kidneys in the same gels lacking RGD approximately maintained their initial midplane area (**Fig. 4F, Fig. S13**). We hypothesized that the inability of kidneys to adhere to the 3D matrix would not only adversely affect organ shape and size but also finer-scale development. Indeed, we discovered a significant 10.3% decrease in nephron:UB tip ratio in kidneys cultured without RGD (p = 0.034, **Fig. 4G**). Taken together, boundary conditions of sufficient stiffness and adhesion are necessary to retain in vivo-like kidney shape and increase in organ size over time. Second, nephrogenesis efficiency (nephron:UB tip ratio) increases for boundary conditions having a ‘goldilocks’ stiffness and higher adhesion (**Fig. 4H**).

## Discussion

We developed a 3D kidney culture technique for the study of morphogenesis dynamics and the effect of matrix material boundary conditions on development. 3D embedding in ECM-derived composite hydrogel rescued *in vivo*-like branching properties as compared to traditional flattened culture, validated with a computational model. Changing the ECM-gel concentration altered the UB tip distribution at the organ surface. By introducing engineered AHA hydrogels that allow independent control of stiffness and adhesion, we discovered that both factors contribute to the overall 3D organ shape as well as nephrogenesis efficiency in culture.

Kidney culture techniques have evolved from metal grids supporting cloth/filter materials supplied with inductive signals from the spinal cord^9,10,66^ to commercially available transwells and other formats imposing ALI conditions^11,12,67^. However, outcomes in ALI culture can differ from those *in vivo*. For example, BMP7 inhibition in ALI culture leads to rare UB tip collision events, while knockout of *BMP7 in vivo* does not affect tip:tip distance^20,68^. Similarly, HGF signaling promotes kidney branching in culture, while *HGF^-/-^* mouse embryos show normal ureteric bud organization^69,70^. These discrepancies motivate improved culture approaches, particularly to avoid gross tissue flattening that occurs in ALI culture. We present a 3D culture approach that better recapitulates *in vivo* morphogenesis. Beyond this study, it will be informative to reexamine known ALI culture artifacts in the 3D format and to revisit mouse models of kidney defects, many of which are poorly understood at the level of live cell-level behavior.

Our previous work showed that without an appropriately timed transition in UB tip orientation, tip crowding and repulsion at the organ surface cause a ‘buried tips’ phenotype, where some tips are displaced to deeper tissue layers^18^. In this study, we observed rare buried tip-cap mesenchyme niches in C+M gels but not in AHA gels (**Fig. S14**). Buried tips are predicted to occur when the available organ surface area per niche falls below a minimal threshold. In the case of C+M gels, the shift in the shape of kidneys from oblong to spherical would reduce their surface area:volume ratio, consistent with the emergence of buried tips. A similar logic follows for ALI culture, where buried tips (relative to the strip of circumferential organ area) are common. This underscores the importance of appropriate control over global organ shape for the kidney to maintain correct positioning of UB tips at the surface during branching.

Our data show that stiffness and adhesion boundary conditions both affect nephron endowment per UB tip in explants. The data complement reports that stiffness and viscoelasticity affects stem cell-derived nephron-lineage kidney organoid differentiation^23–25^. However, the organization of cells in organoids do not closely resemble that of UB and nephron lineages *in vivo*, nor their reciprocal signaling when co-cultured^71–73^. This suggests a cell intrinsic component to interpretation of the mechanical microenvironment. Further work is needed to understand other potential effects of mechanical boundary conditions, for example in setting the length and time scales over which endogenous stresses associated with branching morphogenesis or nephron condensation persist^19,74^. The connection between these factors and nephron progenitor renewal vs. differentiation balance remains an open question.

One area warranting future investigation are alternative materials for 3D embedding that could confer broader engineering control over physical properties and bioorthogonality. Systems with higher or lower resemblance to the native ECM properties of the kidney microenvironment are open to further study, as well as the effect of viscoelasticity^13,23^. While we focused on hyaluronic acid hydrogels for their biocompatibility and potential for controlled mechanical modification, HA itself is a biologically active ECM component in mesoderm-derived tissues including the kidney. During kidney development, HA is mainly deposited by nephron progenitors around the UB tips^75^. HA regulates both UB branching and nephrogenesis in a concentration and molecular weight-dependent manner when supplied in media^76,77^. Though our HA gels are crosslinked and do not penetrate UB tip niches, unreacted HA chains or those liberated by crosslink degradation could diffuse into kidney tissue. However, we expect minimal confounding effects since the HA used to synthesize our gels has a molecular weight of 70 kDa, similar to a 64 kDa soluble HA condition that previously yielded comparable phenotypes to untreated controls in ALI culture^77^. However, to better isolate the effect of matrix stiffness/adhesion, biologically inert materials such as poly(ethylene glycol) (PEG)-based hydrogels are potential alternatives^78^.

Our work motivates continued investigation of the role of the force balance at the kidney periphery to its global and local organization. Appropriate patterning of tensile, shear, and pressure forces may contribute to new strategies for rational control over morphogenetic modules such as tubule elongation, branching, and cell fluxes in the ureteric bud, cap mesenchyme, and stroma. Such a capability would address engineering barriers to setting nephron condensation, connectivity with the ureteric bud, and corticomedullary patterning toward synthetic kidney replacement tissues.

## Supporting information

Movie S1: E13 kidney in 3D 0.5C+M culture with EPCAM and PNA-lectin over 48 hrs.

Movie S2: Movies of tip movement tracking in ALI culture over 11 hrs.

Movie S3: Movies of tip movement tracking in 0.5C+M culture over 11 hrs.

Movie S4: GDNF upregulation in 3D culture over 21 hrs.

Movie S5: Movie showing the effects of RET downregulation over 21 hrs.

Movie S6: Embryonic kidney exerts contractile force on surrounding labeled collagen in C+M culture.

Movie S7: Dynamic stromal cell migration in ALI culture.

Movie S8: Dynamic stromal cell migration in 3D culture.

## Acknowledgements

We thank past and current members of the Hughes lab for their help and discussions, especially J. Viola, J. Liu, and S. Grindel. We also thank Y. Zhang and D. Huang for advice on biomaterials, and X. Luo for help with the Rhino model. We are grateful to P. Mollenkopf and P. Janmey for training and assistance with material characterization. We thank G. Timin and M. Milinkovitch for sending image files from ref. ^33^. This work was supported by the trainee pilot award from Center for Engineering MechanoBiology (CEMB), an NSF Science and Technology Center, under grant agreement CMMI: 15-48571 (A.Z.H), NIH F32 fellowship DK126385 and Penn Center for Soft & Living Matter fellowship (L.S.P.), and NSF CAREER award 2047271 (A.J.H.).

## Materials & Methods

### Mouse strain and microdissection

All mouse experiments followed National Institutes of Health (NIH) guidelines and were approved by the Institutional Animal Care and Use Committee (IACUC) of the University of Pennsylvania. Pregnant CD1 mice were purchased from Charles River Production at the desired embryonic ages. Embryo age was confirmed by limb staging^79^. Kidneys as well as whole urogenital tissues were microdissected from mouse embryos and kept on ice before experiments.

### Culture plate fabrication

50 ml of 10:1 (base:crosslinker) Polydimethylsiloxane (PDMS) elastomer (Sylgard 184) was cast onto a 15 cm petri dish and baked at 60°C overnight to form a sheet. The cured PDMS was punched using a 0.625” hole punch and a 12 mm biopsy punch to create PDMS rings. The PDMS was then plasma-bonded to glass-bottom 12-well plates (P12-1.5H-N, Cellvis) in a plasma oven (HPT100, Princeton Scientific) and baked at 60°C for >1 hour to enhance bonding. The plates were sterilized before use and the outer space between the PDMS and the well was filled with PBS with 1X Pen-Strep.

### Live antibody labeling

To live label the explants, kidneys were incubated at 37°C in fluorophore-conjugated antibodies for 2 hours and washed once before culture. Live label antibodies were CD326 with FITC (11-5791-82, Invitrogen) and Lectin PNA From Arachis hypogaea with Alexa Fluor 647 (L32640, Invitrogen) with 1:250 dilution in culture media. In long-term time-lapse experiments, cultured explants were labeled again by adding fluorescent antibodies in media for 1 hour in the incubator and washed before resuming imaging.

### Explant culture in 3D hydrogel and ALI

3D culture: Dissected kidneys were washed in PBS and transferred to PDMS rings with a pre-cut gelatin-coated p20 pipette and excess liquid was removed. 10 ul of hydrogel solution was added to create a droplet and suspend the kidney. The culture plate was incubated at 37°C for 30 minutes for gelation. After gelling, DMEM media with 10% fetal bovine serum (FBS, MT35-010-CV, Corning) and 1x pen/strep (100 IU/mL penicillin, and 100 μg/mL streptomycin, 100x stock, 15140122, Invitrogen) was added to the PDMS ring. ALI culture: ALI culture experiments were primarily done in transwells (3460, Corning). by placing the kidneys on top of a filter membrane with 0.4 µm pores and supplying culture media underneath. The time-lapse movies at ALI were acquired using low-volume dishes modified from Sebinger et al.. Briefly, kidneys were placed in the middle of the PDMS rings and a low volume of media (∼80 ul) was added to generate the air-liquid interface.

### Collagen and Matrigel composite hydrogel

Collagen was prepared as described by Viola & Porter et al.^80^. Rat tail collagen I (354236, Corning, 354236) was neutralized on ice with 1M NaOH, 10X PBS and DI water to the desired concentration. For collagen labeling, NHS-ester(Succinimidyl Ester) with Alexa 555 (A20009, Invitrogen) was added to the collagen stock at 1:1000 the day before. Collagen was neutralized with 1 part labeled collagen and 2 parts unlabeled collagen. human-ESC qualified Matrigel (354277 lot 17823003, Corning) was also labeled using the same technique. Neutralized collagen at 1 mg/ml or 3.6 mg/ml and Matrigel are then mixed at 1:1 ratio to make the 0.5 and 1.8 C+M composite gels, respectively.

### Tip rotation model

The tip rotation model was based on a physics-based form-finding model built in Rhino (v.7, Robert McNeel & Associates) using the Grasshopper Kangaroo2 package (Daniel Piker). It models trees of elastic edges with self-repelling branch nodes enveloped within an elastic bounding domain. Quantitative principles of this software environment are detailed in ref.^18^ We used the SphereCollide goal to model node mutual repulsion at a radius of 80 µm (mean tip-tip distance for E14 kidneys from **Fig. 2B**). We used the Length goal to model tree edge elasticity. Branching nodes below either the grandparent generation (3D) or the parent generation (ALI) were fixed to prevent unrealistic global rotation of tree families as seen in our time-lapse movies. We ran simulated trees within a bounding box of fixed volume at different heights to mimic the 3D and ALI environments. We estimated the initial tree (three sub-trees with two generations of branches) and bounding box geometries from our culture data shown in **Fig. 1D** and **Fig. 2**. The model was run to equilibrium before adding new tips in the same direction as parents using a customized Matlab script. Two new tips were assigned to each terminal end with a 60-degree bifurcation angle either 90 degrees rotated from their parent plane (3D) or along the direction of their parents in the xy plane (ALI). The 3D bounding box was also expanded assuming equal increase in x, y, and z directions based on spherical growth in our 3D culture. The model was run again and 𝜙 and θ angles were measured both before and after this step. Detailed model parameters are included in **Fig. S3**.

### AHA hydrogel

Synthesis of acrylated hyaluronic acid was based on previous methods^60^. Briefly, AHA modification was performed through the reaction of acrylic anhydride (Sigma Aldrich) with sodium hyaluronate (HA, MW = 70 kDa, Lifecore Biomedical) at a pH 9–10 for 3 h at RT in DI water. AHA was purified via dialysis, lyophilized and modification was confirmed using ^1^H NMR (Bruker Neo 400). AHA of 20% modification was confirmed by ^1^H NMR, and used for all experiments (**Fig. S10**, analysis performed in TopSpin). To form hydrogels, RGD(GCGYGRGDSPG, Genescript) and MMP-degradable peptides (GCNSVPMSMRGGSNCG, Genescript) were conjugated to AHA via Michael addition between thiols on cysteines and acrylates during gelation. The reaction was carried out at 37°C for 30 mins in pH 8.5 Media 199 (11150067, Gibco) with 10% FBS and 1% Penstrep. 3% w/v AHA was used for all studies. For the varying crosslinker experiments, all AHAs were functionalized with 1 mM RGD.

### Hydrogel microindentation

Hydrogel preparation: 100 ul hydrogel solutions were added to PDMS molds and incubated at 37°C for 30 minutes to generate *r* = 4 mm cylinders. To mimic the osmotic environment in culture, formed hydrogels are then swelled in culture media overnight in the incubator before taking measurements. AHAs with varying crosslinker concentrations were all fabricated without RGD for microindentation, except for the with and without RGD comparison. Microindentation: microindentation was performed as described by Prahl, Liu, Viola *et al.*^19^. The system consists of a stepper motor attached to a μN-resolution tensiometer that includes a 255 µm cylindrical 30 gauge (AWG) SAE 316L stainless steel wire and a microbalance^81^. The indenter was lowered at a rate of 12.5 µm/s to indent the sample while force, time, and displacement data were recorded. The force-displacement relationship was found to behave as a Hookean spring with force linearly related to displacement, and the spring constant was calibrated using a glass substrate before each measurement.

### 2D AHA cell adhesion assay

3% AHA solution with 2.25 mM MMP-degradable crosslinker and with or without 1 mM RGD peptides was added to 96-well glass-bottom plates and crosslinked as described above. MDCK-II epithelial cells (female, 00062107-1VL, Millipore Sigma) expressing Lifeact-GFP and H2B-mRuby were lifted with 0.25% trypsin-EDTA (2530056, Corning) and plated on AHA with and without RGD at the same density. The MDCKs on AHA were cultured in minimum essential medium (MEM, Earle’s salts and L-glutamine, MT10-010-CV, Corning) supplemented with 10% FBS and 1x pen/strep overnight. Cell adhesion was quantified on the second day by manually annotating attached cells and floating cells in each well.

### Immunofluorescence and optical clearing

Immunofluorescence staining and imaging was performed as previously described^18^, adapted from Combes *et al.* and O’Brien *et al.* ^82,83^. Kidney explants were fixed in 4% paraformaldehyde (J19943.K2, Thermo Scientific), washed twice in 1x DPBS-glycine and once in DPBS. Blocking solution was made by adding 10% donkey serum to an IF wash solution composed of 1 mg/ml bovine serum albumin, 0.2% (v/v) Triton X-100, 0.041%(v/v) Tween-20 in 1X DPBS. Fixed samples were blocked overnight and then incubated in primary antibody in the same blocking solution at 4°C for >2 days depending on their age. A full list of antibodies is provided in **Table 1**. Samples were washed in three exchanges of 1X PBS at least 1 hour each and then incubated with secondary antibodies in blocking solution at 4°C, all with 1:300 dilution in 10% donkey-serum. Stained samples were then cleared using for 2 days in ScaleA2 (4 M urea + 0.1% Triton X-100 + 10% glycerol) followed by 2 days in ScaleB4 (8 M urea + 0.1% Triton X-100)^84^, and imaged in ScaleA2.

**Table 1:**
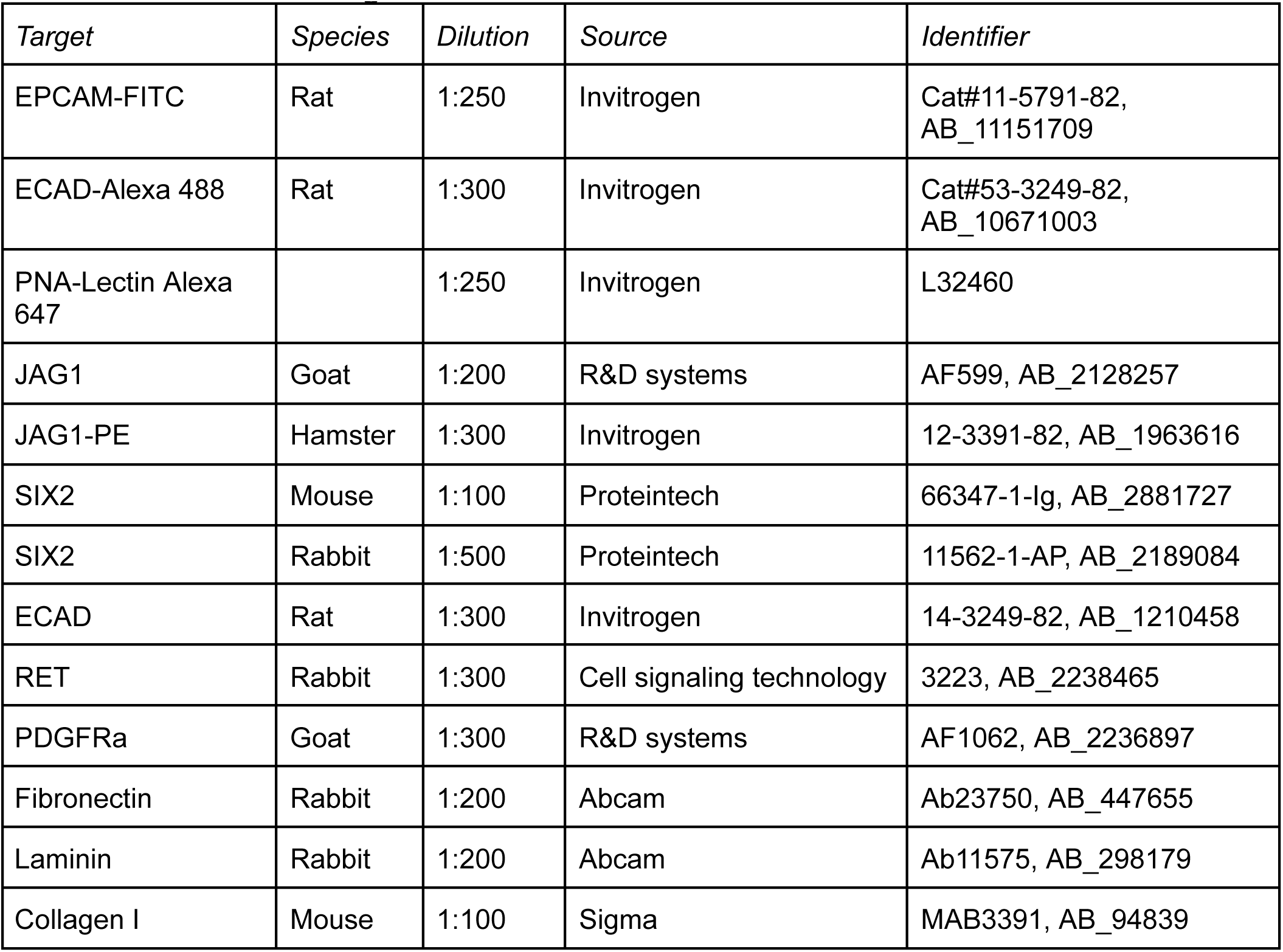
Antibodies and reagents.

### Microscopy

Confocal imaging was performed using a Nikon Ti2-E microscope equipped with a CSU-W1 spinning disk (Yokogawa), a white light LED, laser illumination (100 mW 405, 488, and 561 nm lasers and a 75 mW 640 nm laser), a Prime BSI sCMOS camera (Photometrics), motorized stage, 4x/0.2 NA and 10x/0.25 NA (Nikon), and a humidified stage top environmental chamber (OkoLabs). For time-lapse imaging, xy-positions were defined in the Nikon Elements software and stacks were collected using 15 µm step size every hour with the 10x objective lens. To image the cleared samples, kidneys were immersed in ScaleA2 solution in glass-bottom plates and imaged using either the 4x or 10x lens.

### Kidney morphology analysis

Tip rotation analysis: Time-lapse movies were processed to remove background and speckles and correct for photobleaching over time in FIJI. Maximum projection of the live EPCAM channel was used to annotate tip movements with TrackMate^85^. The segmented tips and tracks were manually filtered to remove misidentified objects. The mean directional change rate was computed by calculating the angle between two consecutive track segments (two time frames) and averaging the angles over the entire track. Kidney structure annotation: UB tips and nephrons were manually annotated from confocal stacks in FIJI using the point tool. UB tips were identified as EPCAM/ECAD+ terminal tips in live samples and RET+ tips in stained samples. Early nephrons were identified by JAG1 expression. Nephron per tip ratio was calculated for each kidney. UB tip-tip distance: For tip-tip distance analysis, neighbors were defined as RET+ UB tips that are at least two generations away (not sharing the same grandparent node) and presented in the same z-plane. The distance was obtained by measuring the distance between the lumen of one tip to its neighbor’s. Kidney cross-sectional shape analysis: The shapes of cultured kidneys were segmented from the maximum projection of the brightfield channel from confocal z-stacks. The outlines were manually traced and measured using the freehand tool and the shape descriptors in FIJI and smoothed by 10% for visual presentation in Fig. 3 and 4. The measurements are defined as: Aspect ratio: major_axis/minor_axis based on the ellipsoid fit of the segmented organ outline. 1 indicates a perfect circle. % area increase: (final area-initial area)*100%/initial area. Initial area was obtained on day 1 and final area was obtained on day 3. Stroma migration area: The stroma migration outlined was obtained by thresholding the brightfield images. The migration area was calculated by subtracting the organ outline area from the total migration outline.

### Statistical analysis

Statistical details for each experiment, e.g., sample size (n), litters, and type of statistical test can be found in the figure legends. Normality test was performed on all datasets before statistical analysis in Prism 10 (GraphPad). Outliers were removed using nonlinear regression (ROUT) method with false discovery rate at 1%. Unpaired two-tailed t-test was used for all comparisons between two conditions. One-way ANOVA with correction for multiple comparisons using Tukley’s test was used for all analyses for three or more conditions within one variable. Two-way ANOVA with correction for multiple comparisons within one variable was used for tests with two variables. Statistical significance denoted as *p < 0.05, **p < 0.01, ***p < 0.001, ****p < 0.0001.

## Supplemental figures

**Fig. S1:**
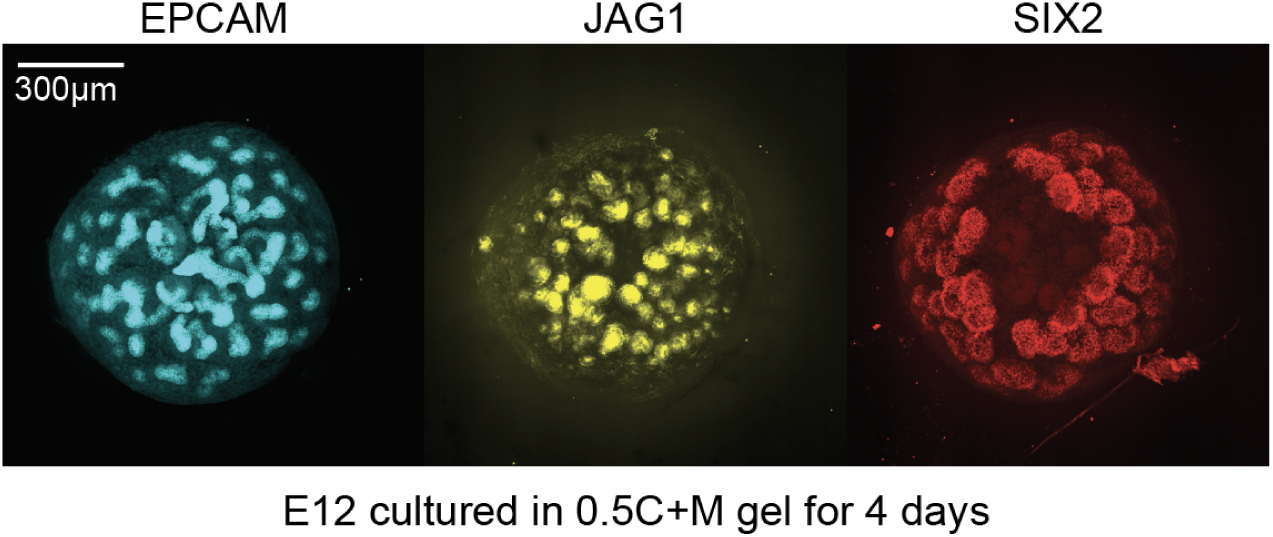
Kidneys develop for at least 4 days in 3D 0.5C+M culture. Immunofluorescence images of E12 kidney cultured in 0.5C+M gel for 4 days, showing markers for the UB and nephron epithelium (EPCAM), early nephron (JAG1), and nephron progenitors (SIX2).

**Fig. S2:**
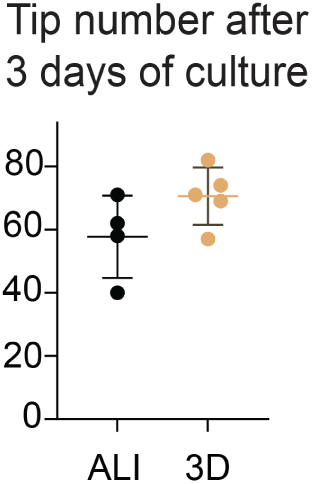
Final tip number is comparable in ALI and 3D culture. Similar data as in **Fig. 1E** where quantification of final UB tip number was annotated from immunofluorescence stacks after 3-day culture in ALI and 3D (0.5C+M) formats, mean ± S.D.

**Fig. S3:**
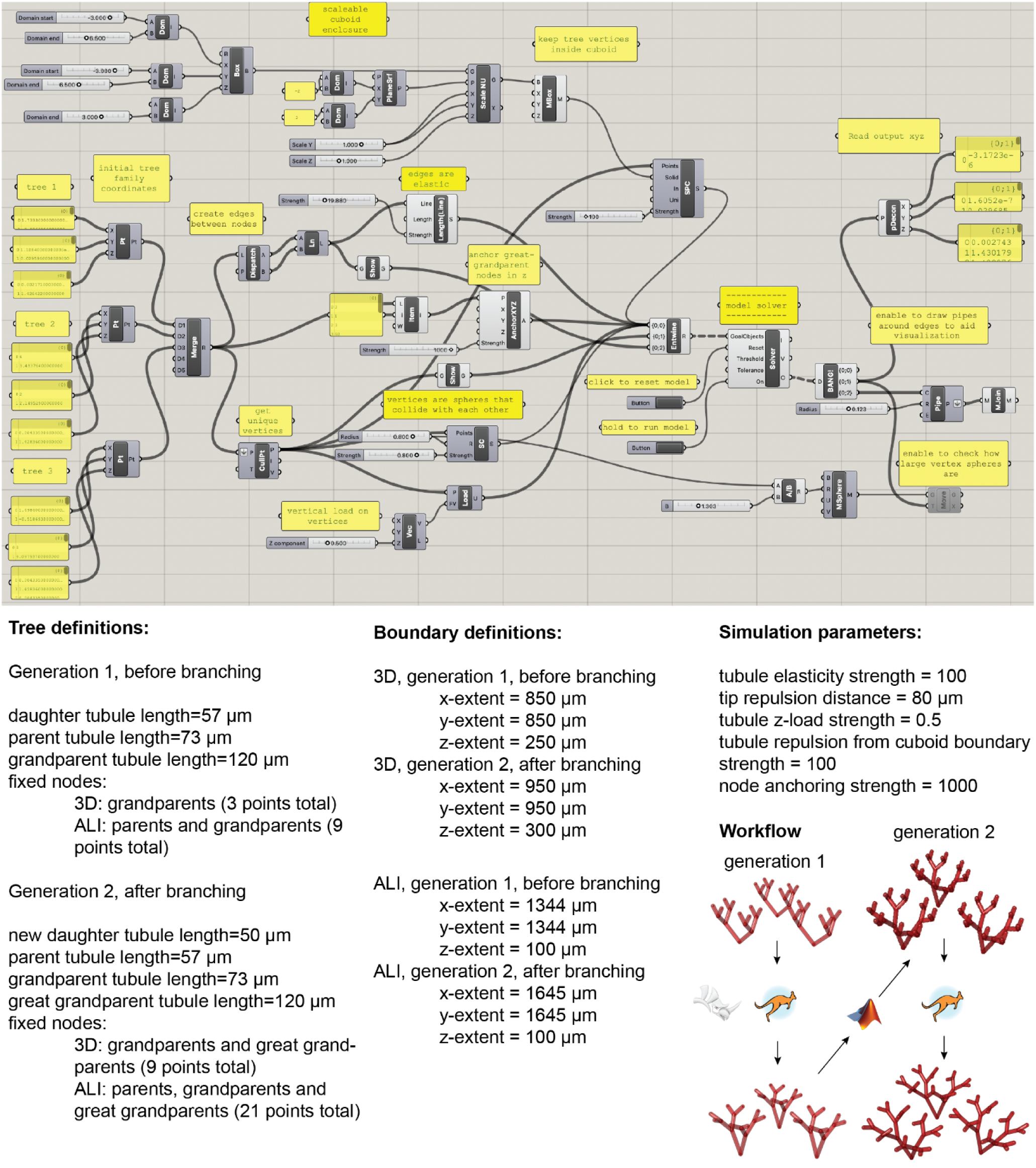
Physics-based model definition. *Top*, Flow diagram of model setup in Rhino Grasshopper. *Bottom*, list of model initial conditions and parameters. *Bottom right*, schematic of model steps, consisting of initial tree setup, energy minimization via the Kangaroo solver, addition of new daughter branches, and a second solver round. Model details are described in main text and **Materials and Methods**.

**Fig. S4:**
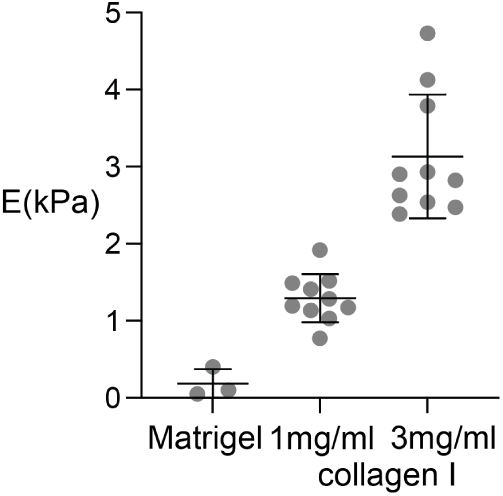
Matrigel and collagen I stiffnesses. Mechanical characterization of each component in the C+M gels by microindentation. Black points are one measurement per gel sample, mean ± S.D.

**Fig. S5:**
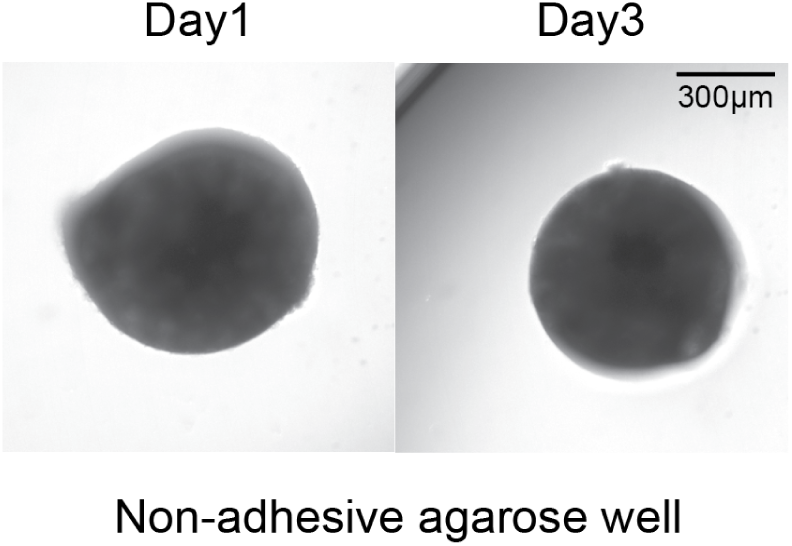
Non-adhesive agarose culture fails to support kidney development. Phase-contrast images of E13 kidneys cultured on 1% agarose-coated plates on day 1 and day 3. Explants fail to develop even under ALI conditions.

**Fig. S6:**
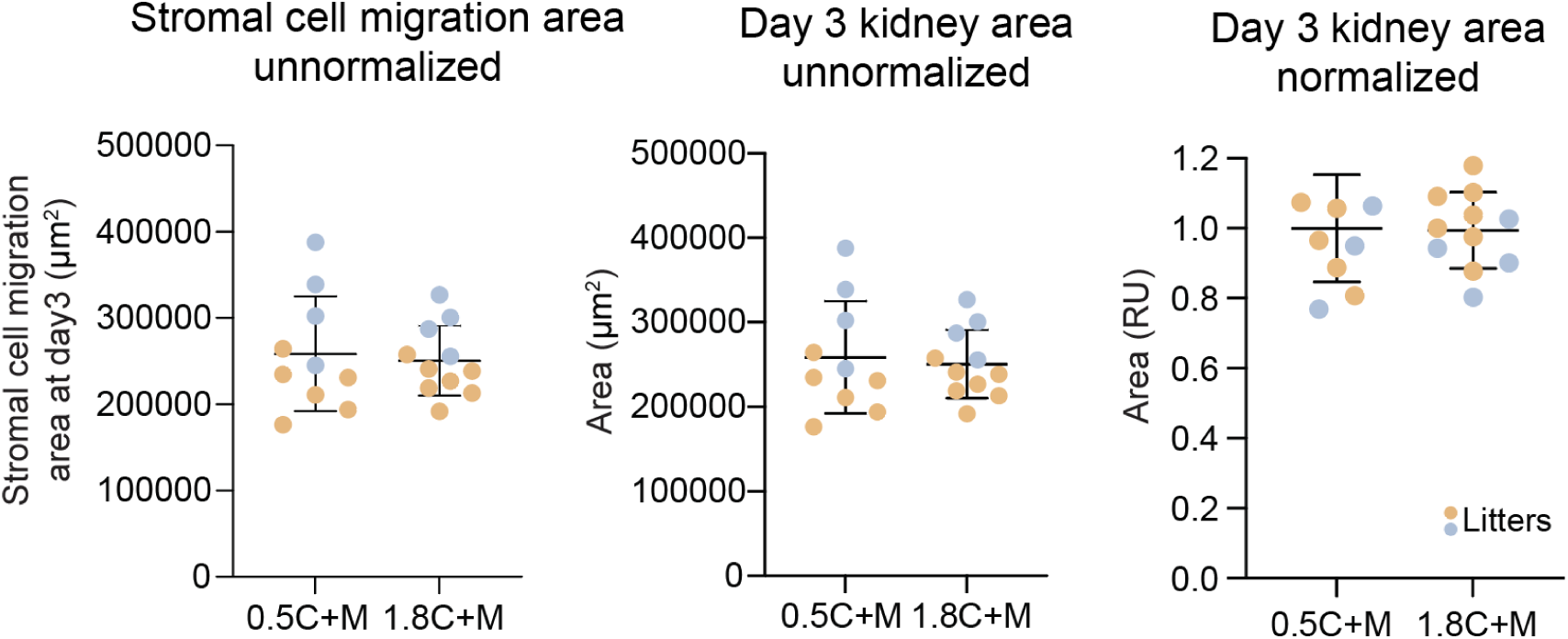
Quantification of stromal cell migration and kidney area in C+M gels. *Left,* unnormalized stromal cell migration area relating to **Fig. 3C**. *Middle-right*, unnormalized, and normalized kidney area. Data shown in the middle plot is used to calculate % area increase in **Fig. 3D**. All plots showing E13 kidney cultured in C+M gels for 3 days.

**Fig. S7:**
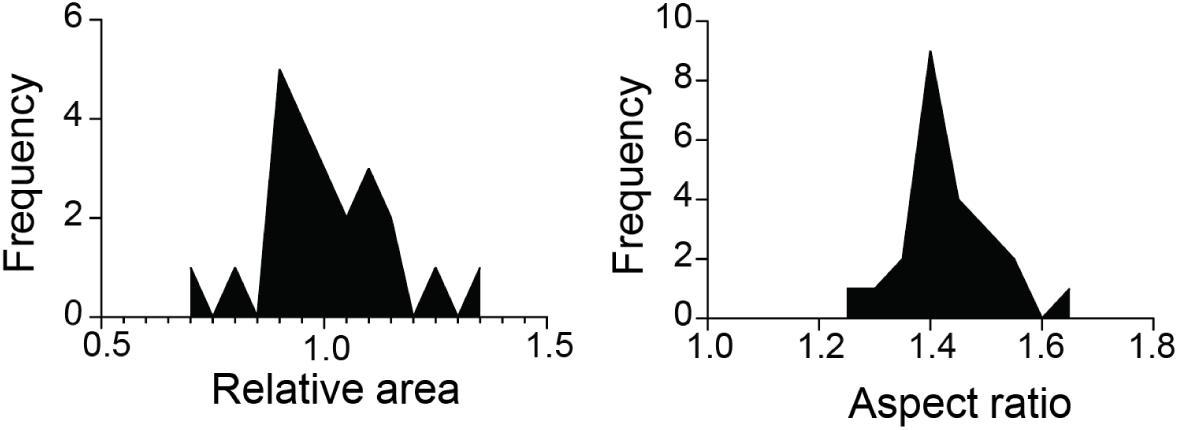
Shape analysis of freshly dissected kidneys showing intra-litter variability. Frequency plots of the relative area and aspect ratio of all 23 kidneys from one E13 litter, showing the within-litter variability in freshly dissected kidneys. Relative area is normalized to the mean area of the litter.

**Fig. S8:**
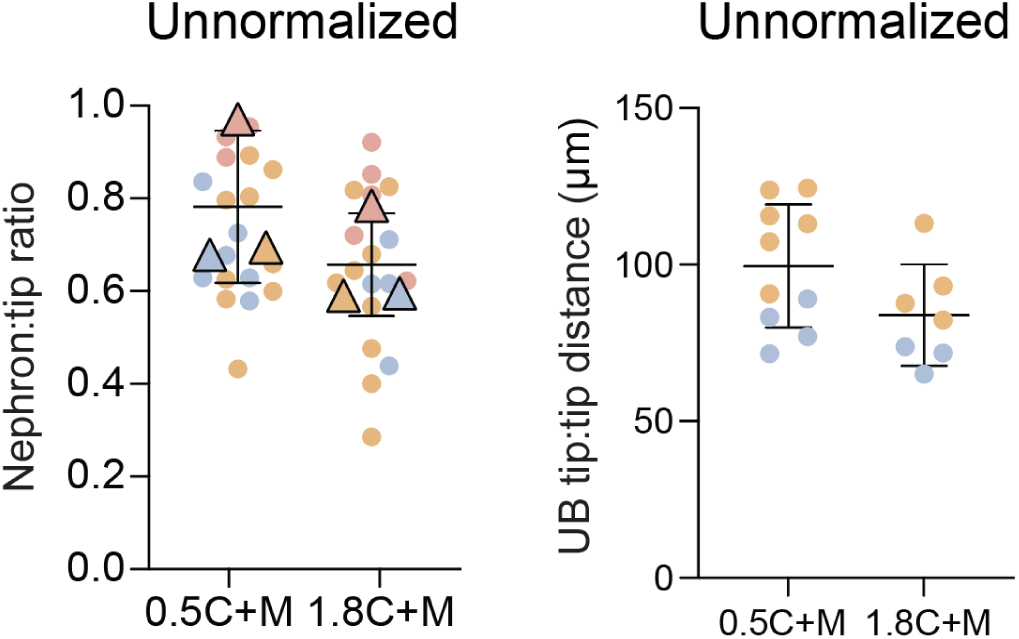
Nephron:tip ratio and UB tip:tip distance before normalization. Plots showing unnormalized data used to generate **Fig. 3E-F**.

**Fig. S9:**
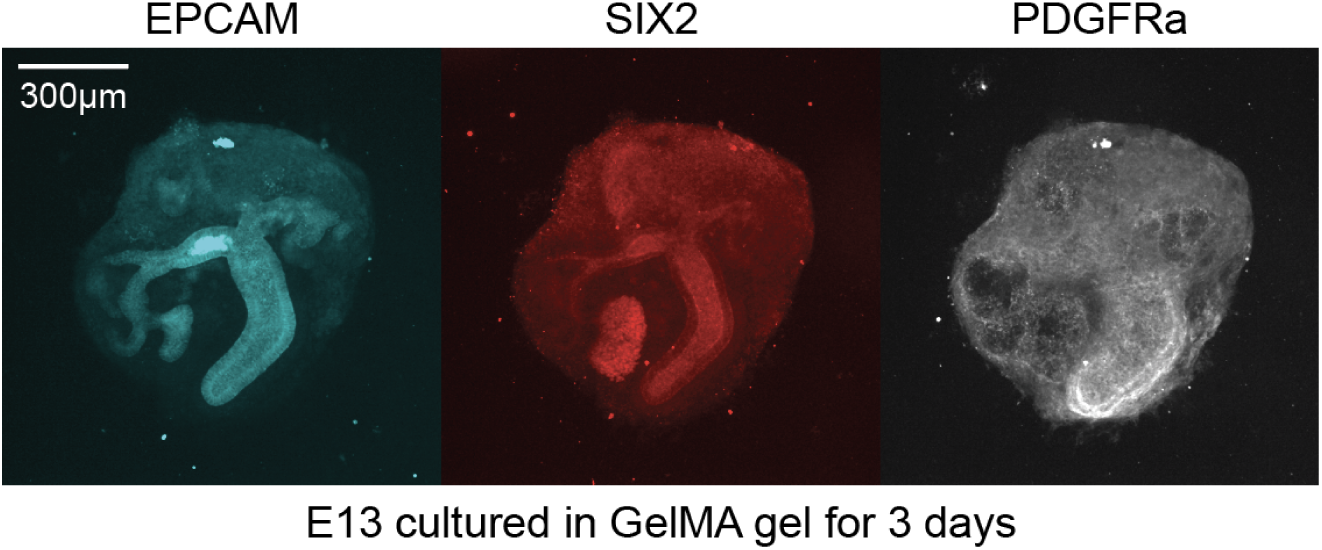
GelMA hydrogel fail to support 3D kidney culture. Representative immunofluorescence images of E13 kidney cultured in GelMA hydrogel for 3 days, indicating poor branching of the ureteric bud and low expression of UB (EPCAM), nephron (SIX2), and stroma (PDGFRa) lineage markers.

**Fig. S10:**
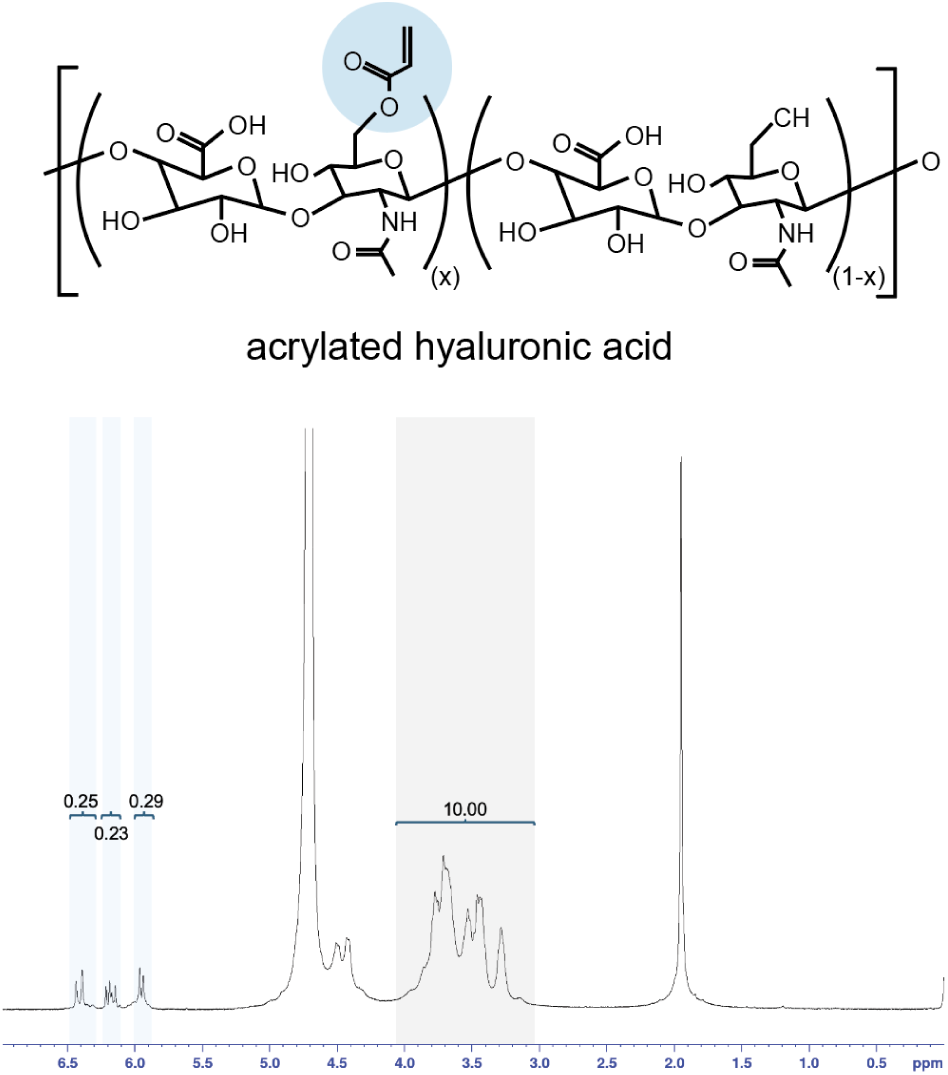
Chemical structure and 1H NMR spectra of synthesized acrylated HA. *Top*, chemical structure of acrylated hyaluronic acid, blue circle indicating the modification of the acrylate group. *Bottom*, 1H NMR spectra showing the integration of the acrylate peaks (Blue, 3H, δ: 5.9-6.1 ppm, 6.1-6.3 ppm, and 6.3-6.5 ppm) normalized to the hyaluronic acid backbone (Grey, 10H, δ: 2.9-4.0 ppm).

**Fig. S11:**
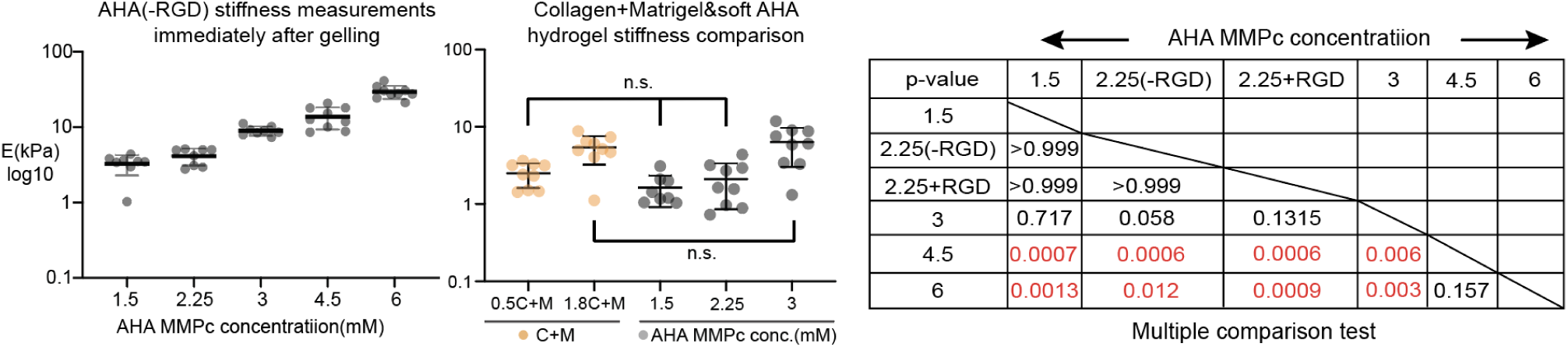
AHA stiffness characterization and statistical results. *Left*: AHA with different crosslinker concentrations immediately after gelling. *Middle*: Stiffness comparison between C+M hydrogel and soft AHA gels. Pairs not statistically different in ordinary one-way ANOVA with Tukey test are indicated as n.s.. *Right*: full statistical results for AHA stiffness after swelling in Figure 4B. Brown Forsythe and Welch Anova test for samples with unequal variance. Significant p-values highlighted in red.

**Fig. S12:**
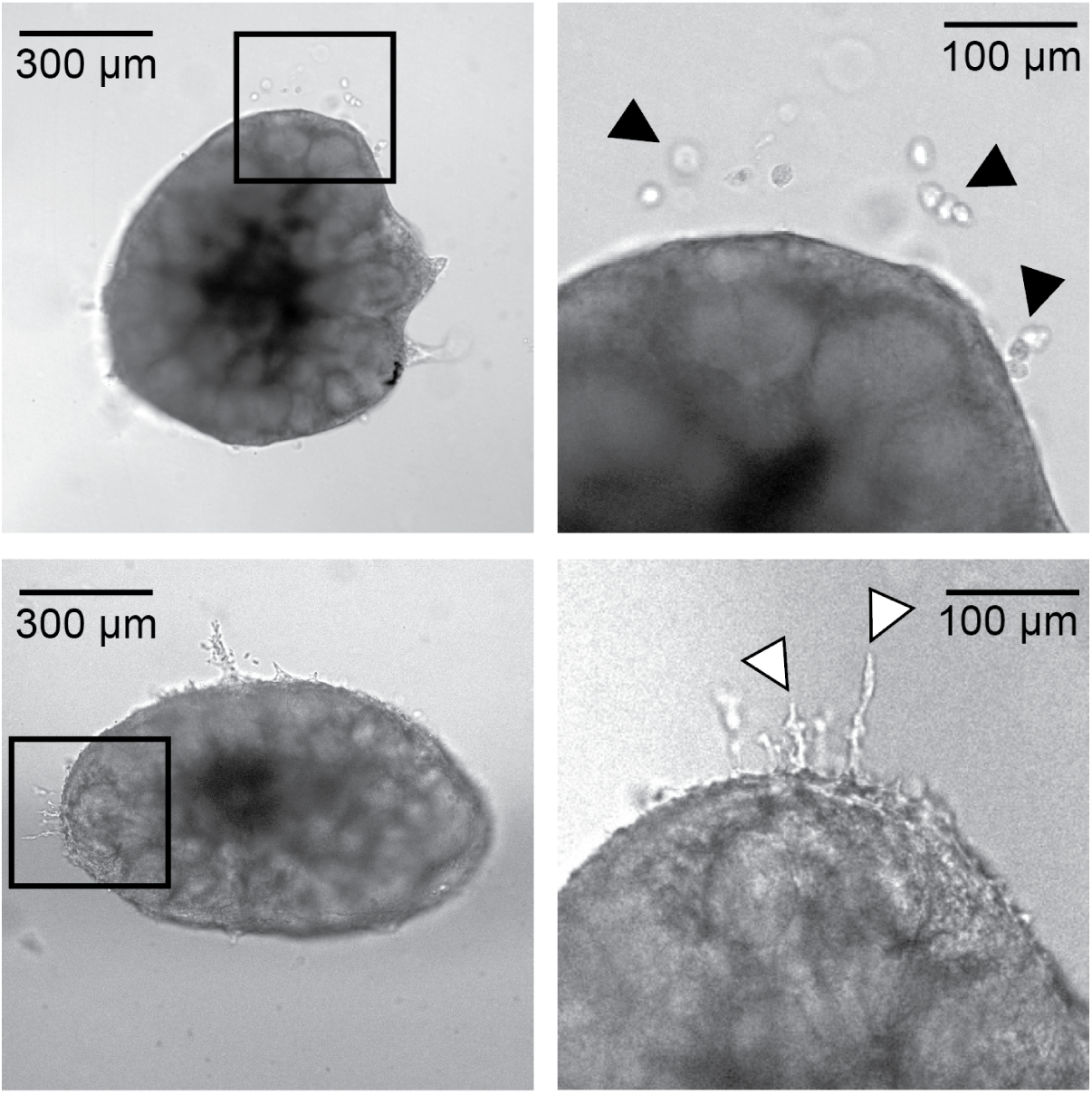
Kidney stromal cells do not adhere to AHA hydrogels without RGD functionalization. Brightfield image and inset showing stromal cells bordering an E13 kidney embedded in AHA hydrogel lacking RGD (*top*) and with RGD (*bottom*) after 3 days in culture. Black arrowheads indicate cells that form non-adhered spheres in the matrix. Black arrowheads indicate cells attached to AHA+RGD.

**Fig. S13:**
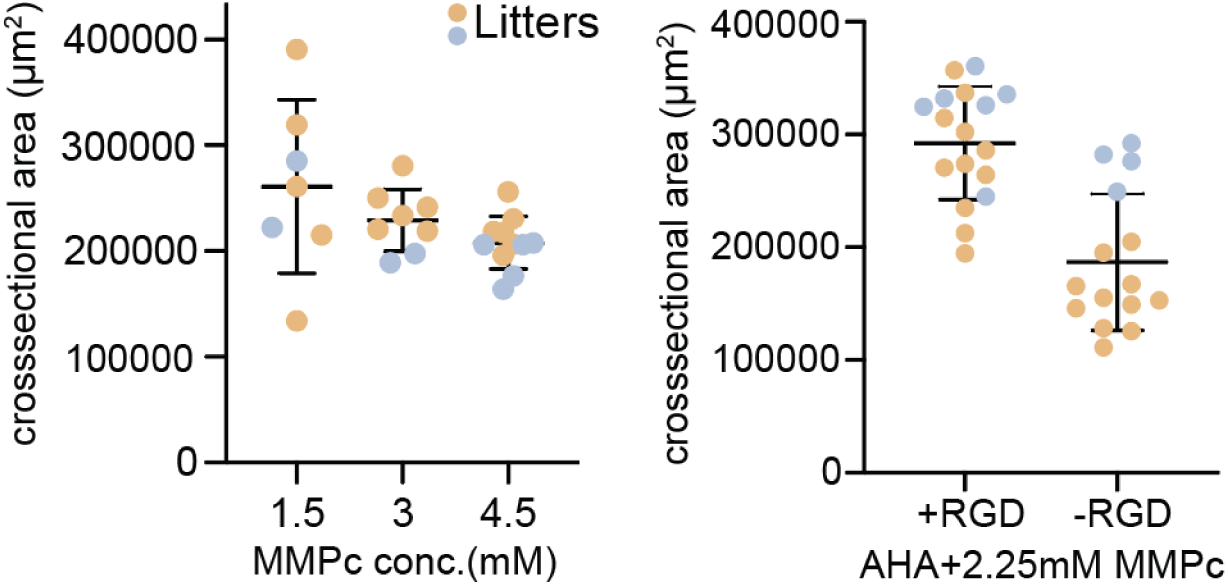
Kidney cross sectional area dependence on stiffness and presence of RGD adhesion ligands in AHA hydrogels. E13 kidneys cultured in AHA hydrogels with different stiffnesses (*left*) and +/- RGD functionalization (*right*). Cross-sectional area analyzed on day 3 of live culture. Data used to calculate % area increase in **Fig. 4D,F**.

**Fig. S14:**
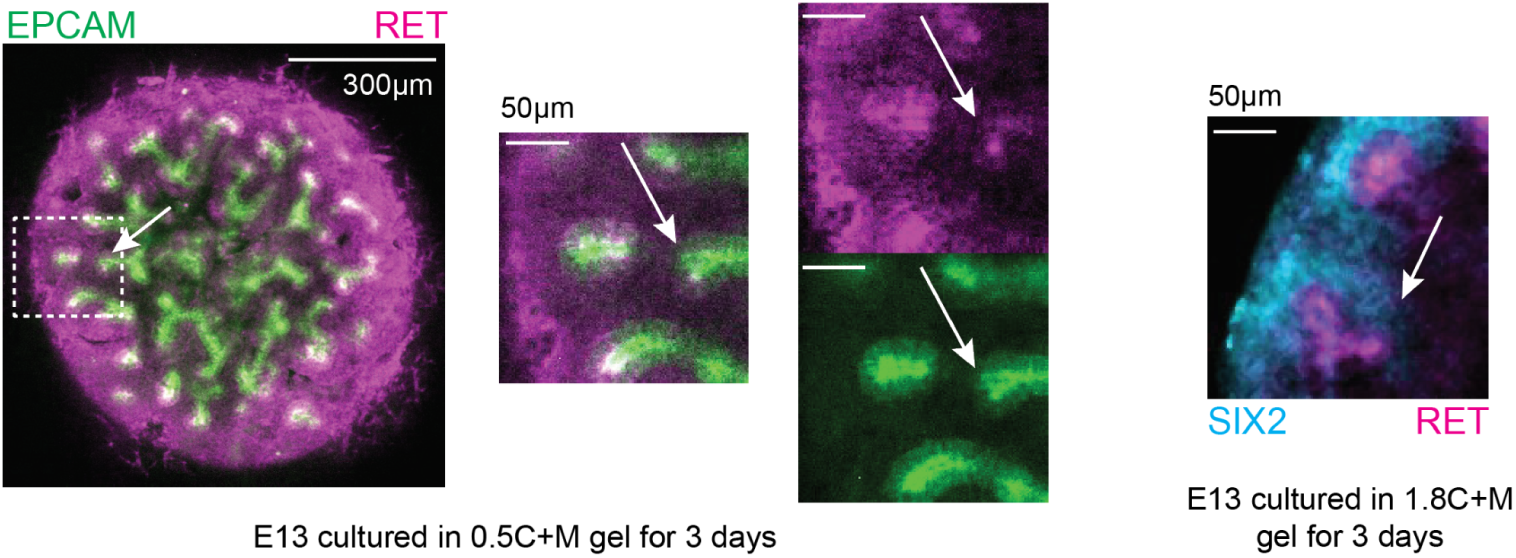
Kidneys cultured in C+M gels acquire buried tips. Immunofluorescence images of E13 kidneys cultured in 0.5C+M (*left*) and 1.8C+M (*right*), both demonstrating examples of buried tips. Arrows indicate tips internalized within the organ, having lower RET expression (*left*) but still surrounded by SIX2+ caps (*right*).

## Supplemental Movies

**Movie S1: E13 kidney in 3D 0.5C+M culture with EPCAM and PNA-lectin over 48 hrs.** EPCAM shown in cyan and PNA-lectin shown in magenta.

**Movie S2: Movies of tip movement tracking in ALI culture over 11 hrs**. Time frames included in Fig. 2D.

**Movie S3: Movies of tip movement tracking in 0.5C+M culture over 11 hrs**. Similar to **Movie S2**, time frames included in **Fig. 2D**.

**Movie S4: GDNF upregulation in 3D culture over 21 hrs.** Time-lapse movies over 19 hrs showing E12 kidney cultured in 0.5 C+M hydrogel supplied with 100 ng/ml recombinant GDNF. Time frames included in **Fig. 2I**.

**Movie S5: Movie showing the effects of RET downregulation over 21 hrs.** *Left*: E13 kidneys cultured in basal media in 1C+M hydrogel. *Middle*: E13 kidney cultured in ALI (transwell) format with 100 nM of the RET inhibitor selpercatinib. *Right*: E13 kidney cultured in 3D 1C+M gel with the same RET inhibition. Time frames included in **Fig. 2K**.

**Movie S6: Embryonic kidney exerts contractile force on surrounding labeled collagen in C+M culture.** Time-lapse movies over 18 hrs showing E13 kidney cultured in prelabled C+M hydrogel pulling fluorescent collagen fibers inward.

**Movie S7: Dynamic stromal cell migration in ALI culture.** Time-lapse movies over 16 hrs depicting periphery stromal cells migrating away from the kidney during ALI culture.

**Movie S8: Dynamic stromal cell migration in 3D culture.** Similar to **Movie S7**, showing stromal cells migrating away from the kidney in 3D culture.

